# Modeling concentration-dependent phase separation processes involving peptides and RNA via residue-based coarse-graining

**DOI:** 10.1101/2022.08.19.504518

**Authors:** Gilberto Valdes-Garcia, Lim Heo, Lisa J. Lapidus, Michael Feig

## Abstract

Biomolecular condensation, especially liquid-liquid phase separation, is an important physical process with relevance for a number of different aspects of biological functions. Key questions of what drives such condensation, especially in terms of molecular composition, can be addressed via computer simulations, but the development of computationally efficient, yet physically realistic models has been challenging. Here, the coarse-grained model COCOMO is introduced that balances the polymer behavior of peptides and RNA chains with their propensity to phase separate as a function of composition and concentration. COCOMO is a residue-based model that combines bonded terms with short- and long-range terms, including a Debye-Hückel solvation term. The model is highly predictive of experimental data on phase-separating model systems. It is also computationally efficient and can reach the spatial and temporal scales on which biomolecular condensation is observed with moderate computational resources.

## INTRODUCTION

Liquid-liquid phase separation (LLPS) is gaining importance in understanding membrane-less subcellular organization. In the cell, liquid condensation is mediated by polymers, mainly proteins and RNA^1^. The first membrane-less compartment observed was within the nucleus of neuronal cells in 1830s, later termed the nucleolus^2^. Today many other compartments that are not delimited by membranes are also known. Examples include the Cajal bodies^3-5^, PML bodies^3^, and nuclear speckles^6, 7^ in the nucleus, as well as the stress granules^8, 9^, P-bodies^10^, and germ granules^11^ in the cytoplasm. More recent studies indicate that biomolecular condensation may be much more ubiquitous beyond these well-known cellular components^12^. Despite differences in composition, location and function the condensates share similarities in shape, dynamics, and assembly mechanisms^13^.

Many biophysical techniques, including microscopy and structural and compositional analysis have been applied to study phase separation (PS) ^14^. On the theoretical side, analytical approaches based on polymer theories have been applied^15, 16^. Finally, computer simulations have been used to explain interactions that stabilize PS, with the most detailed insight derived from atomistic simulations^17-19^. However, atomistic simulations are challenged^20^ by the significant computational resources required to reach the time scales (µs-ms) and spatial scales (>100 nm) on which LLPS is observed experimentally. Coarse-grained (CG) models are a computationally more efficient alternative ^21, 22^ and they have been used successfully to study PS via simulation ^23^. Earlier studies stem from the colloid field with more limited applicability to specific biological systems^24^. More recently, biology-focused models at different resolution levels have been developed, ranging from models representing proteins/RNA at the molecule level as single particles^25^ to patchy particles^26^, residue based models^23, 27-33^, and higher-resolution models with multiple particles per residues^34-36^.

Residue-based sequence-dependent models have become very popular for studying PS as they combine computational efficiency with an ability to retain key physicochemical features of specific biological systems. Many rely on a hydrophobicity scale (HPS) using an Ashbaugh–Hatch modified Lennard-Jones potential^37^ to describe short-range interactions^23, 27-29, 33^. In some cases the HPS are implemented without further optimization^23^, while other models apply machine learning and Bayesian parameter-learning procedures for optimization^27, 29^. Further HPS optimizations have focused on cation-π interactions given its importance in PS^31, 32^. These CG models have been able to reproduce some experimental data reasonably well. One limitation of the existing models, as will be shown below, is that concentration-dependent PS is not reproduced well, which limits their predictive ability since concentration is a key factor governing phase behavior^38^. Moreover, most models focus on describing only protein systems without a compatible nucleic acid model preventing studies of increasingly important peptide-RNA condensates.

Here, we propose a new residue-based CG model, termed COCOMO (COncentration-dependent COndensation MOdel) to describe PS in peptide-only and peptide-RNA systems. Our goal was to develop a simple, yet accurate model for describing coacervation in systems containing only peptides or mixtures of peptides and RNA in a concentration-dependent manner. The model was designed to minimize the number of necessary parameters to maintain as much general applicability as possible, and it introduces a term to account for solvation effects at the residue level. We demonstrate that our model can accurately reproduce experimental PS data, including in systems that were not included in the parameterization, while maintaining a balanced description of individual polymer properties of peptides and RNA molecules.

## METHODS

### Coarse-grained model

In COCOMO, each residue, either a protein amino acid or an RNA nucleotide, is represented as a single spherical particle. The total interaction energy is given by:

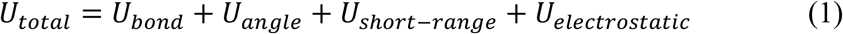

where *U*_*bond*_ is the harmonic potential for chain connectivity:

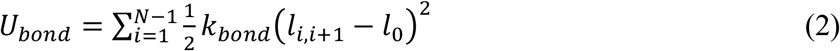

in which *l*_*i,i+*1_ is the distance between two neighboring residues, *k*_*bond*_ is the spring constant, and *l*_0_ the equilibrium bond length. We choose *k*_*bond*_ = 4184 *kJ*/*mol* · *nm*^2^, which is a softer value than in all-atom potentials; for proteins, *l*_0_ = 0.38 *nm*, from the average Cα-Cα distance, and, for nucleotides, *l*_0_ = 0.5 *nm*, corresponding to the average distance between backbones for single-stranded nucleic acids^39^.

*U*_*angle*_ is the angle potential between three neighboring particles to account for chain stiffness:

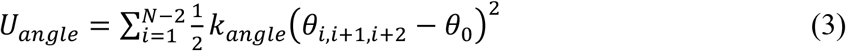

where *θ*_*i,i*+1,*i*+2_ is the angle between three neighboring beads, with angle constant *k*_*angle*_ equal to 4.184 kJ/mol·rad^2^ for proteins, and 5.021 kJ/mol·rad^2^ for nucleic acids. The target angle was set to *θ*_0_ = 180º.

Non-bonded pairwise interactions consist of a short-range 10-5 Lennard-Jones potential, *U*_*short*−*range*_, and a long-range Debye-Hückel potential, *U*_*electrostatic*_, as follows:

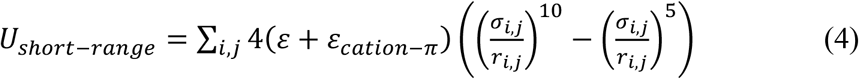

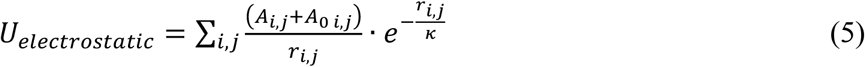

Here *r*_*i,j*_ is the inter-particle distance, *σ*_*i,j*_ is the distance at which the potential is zero, ε is the depth of the potential well, *ε*_*cation*−*π*_ is added to augment cation-π interactions, *A*_*i,j*_ = *A*_*i*_ *∗ A*_*j*_ describes attractive or repulsive long-range interactions, *A*_0 *i,j*_ = *A*_0 *i*_ + *A*_0 *j*_ reflects the effective repulsion between polar residues due to solvation effects, and *κ* is the Debye-Hückel screening length.

Optimized non-bonded parameters values are as follows: *ε*_*polar residues*_ = 0.4 *kJ*/*mol, ε*_*non*−*polar residues*_ = 0.41 *kJ*/*mol*. Arg, Asn, Asp, Cys, Gln, Glu, His, Lys, Ser, and Thr were considered polar residues; Ala, Gly, Ile, Leu, Met, Phe, Pro, Trp, Tyr, and Val were considered non-polar. For nucleotides, we used *ε*_*nucleotides*_ = 0.41 *kJ*/*mol*. We further adjusted for cation-π interactions by adding *ε*_*R*/*K* − *F*/*Y*/*W*_ = 0.3 *kJ*/*mol* for interactions within proteins and *ε*_*R*/*K* − *nucleic*_ = 0.2 *kJ*/*mol* for protein-RNA cation-π interactions. The effective radii 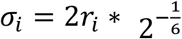 were set from the radius, *r*_*i*_, of a sphere with equivalent volume of a given residue. *A*_*i*_ was calculated from residues charge (*q*_*i*_) +1 for Arg/Lys, -1 Asp/Glu and nucleotides, and 0 for the rest according to 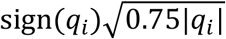 as proposed previously^25^. *A*_0_ was set to 0.05 for polar residues and nucleotides, it was set to 0 for non-polar residues. Finally, *κ* = 1 nm, except when noted, corresponding to an ionic strength of ∼100 mM. **Table S1** reports all residue-specific parameters employed in the model.

### Molecular dynamics simulations

The model was simulated by molecular dynamics simulations using OpenMM 7.7.0^40^. Langevin dynamics was applied with a friction coefficient of 0.01 ps^−1^. Initially, a 5,000 step steepest descent minimization was performed followed by 20,000 steps of MD with a time step of 0.01 ps. After that, systems were run for production using a time step of 0.02 ps. Non-bonded interactions were calculated using periodic boundary conditions and truncated at a cutoff distance of 3 nm. Residues separated by one bond were excluded from non-bonded interactions energy calculation.

Individual protein and RNA chains were simulated in a cubic box with a side length of 300 nm at 298 K. For systems described here, five replicates were run over 500 ns, saving coordinates every 200 ps for each system. This was long enough to establish converged ensembles from which average radii of gyration and persistence length could be extracted. The first 100 ns of each trajectory were excluded from the analysis. An initial random conformation for each chain was obtained using a custom python script. Topology files were generated with the MMTSB Tool Set^41^ and CHARMM v44b2^42^. Using a GeForce RTX 2080 Ti GPU card, we could simulate 100 ns of a 100-residue protein in 5 minutes.

Polymer properties were calculated and averaged from the five replicates. For protein sequences we calculated the radius of gyration using MDTraj library ^43^. For RNA chains we also determined the end-to-end distance and the orientational correlation factor (OCF) as a function of separation along the chain |*i* − *j*|, calculated according to:

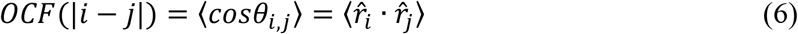

where 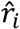 and 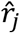 are normalized vectors between any *i, i* + 1 and *j, j* + 1 bonded residues in the chain, respectively.

Systems consisting of multiple chains, either proteins or protein-RNA mixtures, were simulated in boxes of side length ranging from 100 to 200 nm to match different concentrations for different systems (see below). Multiple-chain systems were assembled as follows: (i) short simulations were carried out for each of the system components as described above, (ii) representative conformations were obtained as cluster centers from RMSD-based clustering using the *MiniBatchKMedoids* method implemented in MSMBuilder^44^, (iii) conformations were randomly picked and placed in the simulation box at random positions and with random orientations, but avoiding any two beads between different molecules to be closer than 5 nm, using a custom python script, until the desired concentration was reached, (iv) topology files were generated for the assembled systems using the MMTSB Tool Set^41^ and CHARMM v44b2^42^. For all systems, one replicate was run for 20 µs, with coordinates saved every 500 ps. This was considered long enough to determine the ability to form condensates as typical times for nucleating condensate formation were on the order of microseconds. In the concentrated systems, we evaluated the ability of our model to reproduce heterotypic and homotypic PS in a concentration-dependent manner. In addition, temperature phase diagrams were constructed for some systems by running simulations at fixed initial concentrations at temperatures ranging from 250 to 310 K. Using a GeForce RTX 2080 Ti GPU card, we could simulate 100 ns of a 30,000-residue system in 15 minutes.

Clustering analysis on multichain systems was performed via contact-based criteria. Using in-house python script, we calculated pairwise distances between residues. Two residues were considered in contact if they were closer than the cutoff distance of 0.9 nm, and two chains were considered part of the same cluster if they have at least one contact between any of their residues. We computed the largest cluster size along simulation time and cluster size distribution in terms of the number of members.

To characterize the protein-protein and protein-RNA aggregates formed in different systems we calculated the mass concentration radially from the cluster’s center of mass (COM) to a distance equal to half of the box side length. The results were fitted to a sigmoid curve:

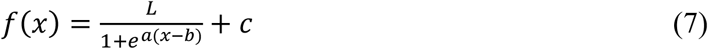

where ***L, a, b***, and ***c*** are fitting parameters. Conveniently, ***b*** is the distance where the concentration drops to half the value from cluster COM and allowed us to determine the cluster dimension. We calculated the density of condensed and diluted phases as the average density at distances ≤ ***b*** /2 and ≥ 2***b***, respectively.

A Jupyter notebook illustrating how to run the model via openMM along with sample analysis is available on github at: https://github.com/feiglab/cocomo

### Model parametrization and test systems

To parametrize the model, we considered a set of 45 intrinsically disordered or unfolded proteins ranging from 8 to 198 residues in length (**Table S2**), primarily to reproduce the experimentally measured radii of gyration (**Table S3**). We note that some of the systems are true disordered peptides while for others disorder was induced by denaturants. A second set of 26 intrinsically disordered proteins not included in the set used for parameterization (**Tables S4-S5**) was used for validation. PolyAde-30, polyUra-30, and polyUra-40 were used to parametrize RNA single chain stiffness (**Table S6**). In addition, parameterization focused on reproducing the concentration-dependent homotypic PS for FUS LCD and LAF-1 RGG peptides (**Table S7**). The model was then validated for predicting PS in three additional homotypic systems (A1 LCD, hTau40-k18, and Ddx4, **Table S7**), for heterotypic protein systems (FUS LCD with FUS RGG3 or [RGRGG]_5_, **Table S8**) and protein-RNA systems for which PS has been reported experimentally (**Table S9**).

## RESULTS AND DISCUSSION

### Model parametrization

The bonded and non-bonded terms in the COCOMO model were parametrized mainly via iterative parameter scans. The main goal was to maximize the agreement with experimental radii of gyration (*R*_*g*_) based on *χ*^2^ values for the IDP systems given in Tables S2 and S3, reproduce RNA polymer parameters given in Table S6, and reproduce PS at the concentrations given from experiment for FUS LCD and LAF-1 RGG peptides.

The bonded terms are comprised of bond and angle potentials. The bond length was based on geometry and a *k*_*bond*_ value “softer” than those from an all-atom potential was chosen. Our chosen value was in the range of other residue-based models published previously^23, 27-29^ and it was not optimized further. For the angle potential, we parametrized *k*_*angle*_ to reproduce *R*_*g*_ distributions for intrinsically disordered proteins (IDPs) (**Figure 1**), and to reproduce *R*_*g*_ and persistence length values for RNA (**Figure 2**). We note that other recently proposed residue-based models for IDPs do not have an angle term^23, 27-29^.

**Figure 1.**
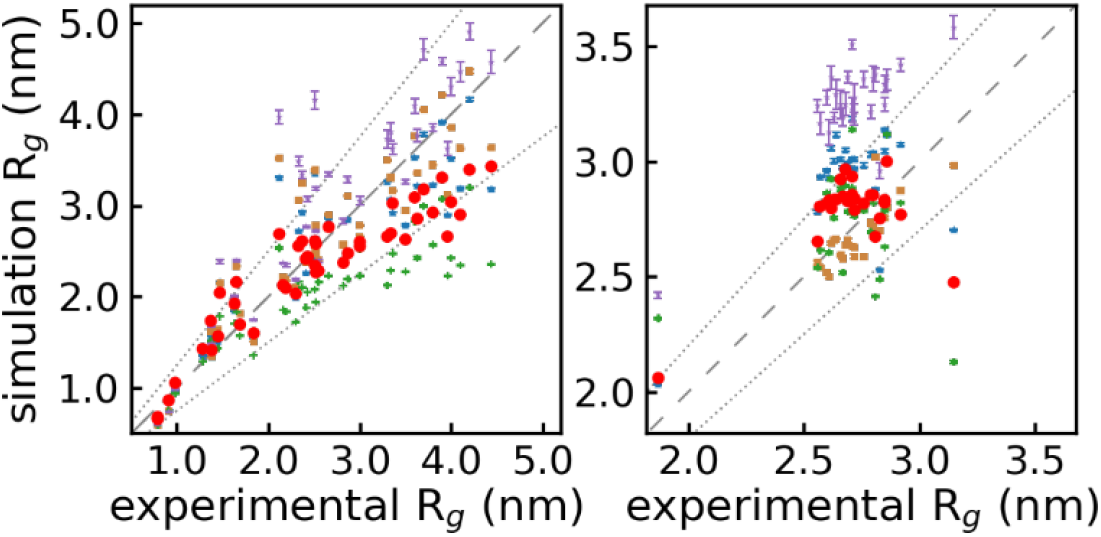
Experimental *vs* simulated radius of gyration. Scatter plots of parametrization set with 45 proteins (A) and the validation set consisting of 26 proteins (B) are shown for COCOMO (red circles) and results obtained by us using the models by Regy *et al*. 2021^33^ (blue triangle), Tesei *et al*. 2021 (tan square), Dignon *et al*. 2018^23^ (green filled plus), Dannenhoffer *et al*. 2021^27^ (purple star). Error bars indicate standard errors from variations between five replicates. Deviations between simulated ensembles and experimental values according to 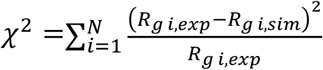 were 3.69, 3.39, 3.50, 7.92, and 6.45 in the parametrization set (A), and 0.40, 0.95, 0.10, 0.81, 3.1 in the validation set (B), respectively for COCOMO and the other models in the order listed above. As a guide to the eye, a dashed line indicates the identity function, and a dotted line shows 25% (A) and 10% (B) deviations from the experimental *R*_*g*_ values.

**Figure 2.**
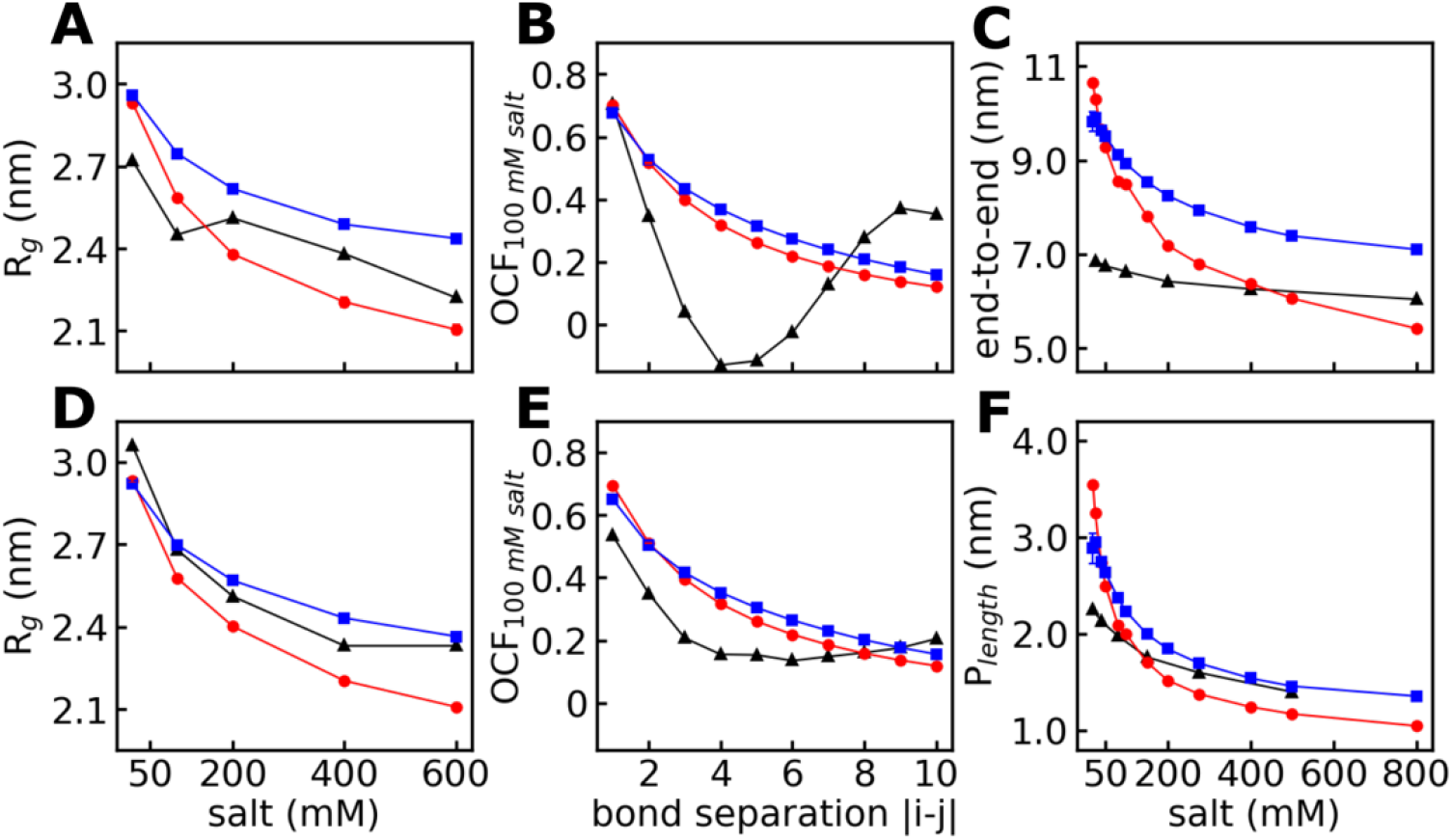
Polymer properties of short RNA sequences. Effect of charge screening on the radius of gyration (*R*_*g*_) of polyAde-30 (A), and polyUra-30 (D), end-to-end distance of polyUra-40 (C), and persistence length of polyUra-40 (F). Orientational correlation factor (OCF) for polyAde-30 (B), and polyU-30 (E) at 100 mM salt concentration is also shown. Results are shown in red circles, blue squares, and black triangles for COCOMO, the Regy *et al*. 2020^33^ model, and experimental values^50, 52^, respectively. Different salt concentrations were reflected by varying the Debye length, *κ*, in Eq. 5 as they can be related according to 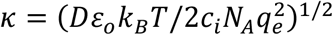, where *D* is the dielectric constant equal to 80, *ε*_*o*_ is the permittivity of free space, *k*_*B*_ is Boltzmann’s constant, *T* is the temperature, *c*_*i*_ is the salt concentration, *N*_*A*_ is Avogadro’s constant, and *q*_*e*_ is the charge of an electron.

Non-bonded interactions affects both polymer properties and PS behavior, so both properties were considered together during optimization. For short-range interactions, we initially started with a single value of *ε* for all interactions, but we found slightly better performance when separating *ε* values between polar and non-polar residues, although the final optimized values that are very similar. It was more important to account explicitly for cation-π interactions that contribute significantly to PS^45-49^ by increasing *ε* values for protein/protein, *ε*_*R*/*K* − *F*/*Y*/*W*_, and protein/RNA, *ε*_*R*/*K* − *nucleic*_, interactions involving interactions between basic amino acids and aromatic moieties. Long-range interactions were mostly determined by nominal charges of amino acids and/or RNA bases, but an additional correction term, *A*_*0*_, was applied to polar residues to effectively account for solvation effects by creating weak repulsion relative to interactions between hydrophobic residues.

The sensitivity of the model to each of the finally chosen parameters is illustrated with the analysis shown in **Figures S1** and **S2**.

### Sampling of intrinsically disordered proteins (IDPs)

The performance of COCOMO on the training (**Table S2)** and validation (**Table S4**) sets in terms of reproducing experimental *R*_*g*_ values is shown in **Figure 1**. For comparison, we also ran individual protein chains with four recently published residue based coarse-grained models^23, 27-29^. The results of our model are similar to the top performing models among those we tried (**Figure 1A**) based on χ^2^ values of 3.69 for the training set and 0.4 for the validation set. For the majority of proteins, the simulated *R*_*g*_ fall within 25% and 10% of the experimental values in the parametrization and test set, respectively (**Figure 1**, dotted lines). The agreement between simulation and experiment is generally good, but as with most other models, our model also shows systematic deviations where smaller systems tend to be less compact, whereas larger systems are more compact than suggested by the experimental values.

To test how *R*_*g*_ is affected by the parameters of our model, we systematically varied one parameter after another while keeping all other values at their final optimized values and repeated simulations. (**Figure S1**). Our results show that variations of *A*_0_, *ε*_*polar*_, and *ε*_*non*−*polar*_ have a strong effect on R_g_, *k*_*angle*_ has a moderate effect, and *ε*_*cation*−*π*_ and *k*_*bond*_ have minimal effects **(Figure S1**). This analysis shows that slightly more optimal parameters could be found if the goal is only to reproduce the experimental *R*_*g*_ values. However, with those values, the concentration-dependent PS behavior in concentrated systems is not reproduced correctly (see below).

### Sampling of RNA

Experimental data on polymer properties is available for polyAde-30, polyUra-30, and polyUra-40. Individual chains of these polynucleotides were simulated using COCOMO and compared with the RNA bead model developed by Regy *et al*. 2020^33^. The results show good agreement with the experimental measurements of *R*_*g*_, end-to-end distances, and persistence lengths (*P*_*length*_) (**Figure 2A, Figure 2C, Figure 2D**, and **Figure 2F**). We also calculated the orientation correlation function (OCF) to quantify the directional persistence of the chain. The correlation decreases with chain distances (**Figure 2B** and **Figure 2E**) similar to experimental values. However, short-range interactions due to base stacking^50^ between bases separated by 3-5 bases that give rise to a dip in the correlation function in the experiment, especially for polyAde-30 are not reproduced because our model does not include beads to represent side chains.

In the simulations a Debye-Hückel term was used to treat electrostatic screening of charge interactions by ions in solution. Therefore, a variation of ionic strength could be modeled by changing the screening length, *κ*. This treatment of electrostatics has been successful in reproducing the ionic strength dependance of polyelectrolyte macromolecules association in similar models^51^. We also find here that the experimental trends are well-reproduced with a decrease of *R*_*g*_ and *P*_*length*_ values at higher salt concentration as in the experiments (**Figure 2A, Figure 2D**, and **Figure 2F**). Best agreement is observed around 100 mM salt concentration (*κ* = 1 nm) which was the condition chosen for our simulations.

Compared to the Regy *et al*. 2020^33^ model, COCOMO agrees similarly well with the experimental data, but while the Regy *et al*. model generates polymers that are slightly less compact than in experiment, COCOMO generates conformations that are slightly more compact.

The sensitivity of RNA simulations to the value of parameters was also evaluated. As observed for proteins, the *R*_*g*_ and *P*_*length*_ values are most sensitive to the choice of *A*_0_ and *ε*_*non*−*polar*_ values, while the choice of *k*_*angle*_ and *k*_*bond*_ showed much smaller effects (**Figure S2**).

### Protein homotypic phase separation

The main focus of COCOMO is on modeling PS phenomena with a model that also maintains realistic polymer behavior of individual molecules. We begin by describing the performance on homotypic systems. We parameterized the model using data from the FUS LCD and LAF-1^RGG^ proteins and then tested it with hTau40-k18 and A1 LCD. Detailed information on the simulated homotypic systems is given in **Table S7** and results are summarized in **Figure 3**. We observe PS dependent on the concentration for all systems in good agreement with experimental thresholds (**Figure 3**, dashed lines)^53-56^. For comparison, we ran the same protein system using previously published modes^23, 27-29^. With those other models, PS was only found for FUS LCD with the Tesei *et al*. 2021^29^ and Dignon *et al*. 2018^23^ models, and even in those cases, PS occurred at all of the concentrations tested here (**Figure 3** and **Figure S3**). No PS was observed with the Regy *et al*. 2021^28^ and Dannenhoffer *et al*. 2021^27^ models for any of the systems tested (**Figure 3** and **Figure S3**). For these two models, we further increased the FUS LCD concentration to 0.4, 0.6, and 0.72 mM. We found that Regy *et al*. 2021^28^ model showed cluster formation at higher concentrations, but with the Dannenhoffer *et al*. 2021^27^ model no PS was observed, even at the higher concentrations (**Figure S4**). This, suggests that the Regy *et al*. 2021^28^ model may be able to at least qualitatively describe concentration-dependent PS, but with the concentration threshold shifted to larger values.

**Figure 3.**
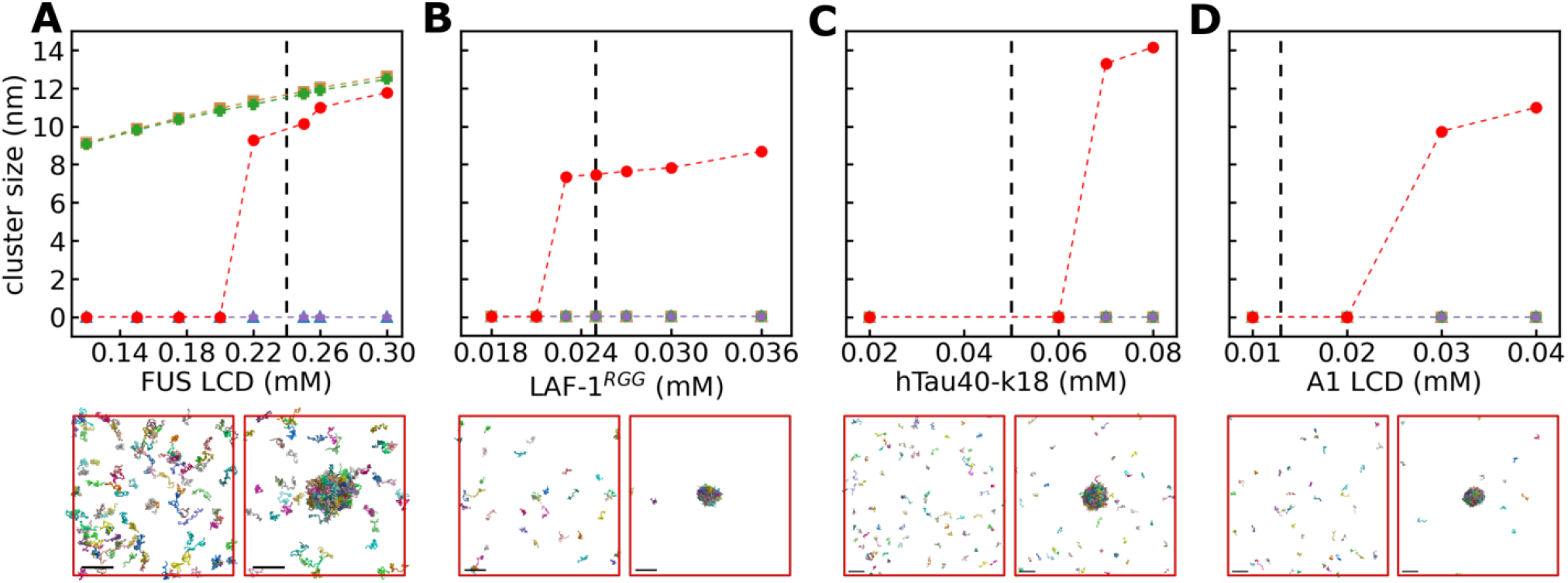
Protein homotypic phase separation. Cluster sizes as a function of concentration and simulation snapshots are shown for FUS LCD (A), LAF-1^RGG^ (B), hTau40-k18 (C), and A1 LCD (D). Cluster sizes were averaged over the last 4 µs of the simulation. Results with COCOMO (red circles) are compared with results using models from Regy *et al*. 2021^28^ (blue triangle), Tesei *et al*. 2021^29^ (tan square), Dignon *et al*. 2018^23^ (green filled plus), and Dannenhoffer *et al*. 2021^27^ (purple star). Experimental LLPS concentration thresholds^53-56^ are shown as dashed lines. The final frames of simulations using our model (lower panels) are shown for the lowest (left) and highest (right) concentrations of each protein system. Coloring is used to indicated different chains. The size bar represents 20 nm.

Analysis of the trajectories showed that the largest cluster size generally stabilizes by 10 µs and increasing concentration accelerates the time necessary to form clusters (**Figure S5** and **Figure S6**). In the case where condensates were not observed, it is in principle possible that condensates did not form due to slow nucleation near the condensation concentration threshold. To test for this possibility, we performed additional simulations for the FUS LCD system starting from the final snapshot at 0.22 mM, with a formed condensate, but then increased the box sizes to lower the concentrations to 0.12 to 0.20 mM, where PS was not observed when starting from randomly distributed polymers. The initial condensate melted at all concentrations except for 0.20 mM (**Figure S7**), indicating that there is only slight hysteresis around the reported concentration thresholds due to slow nucleation kinetics when forming or melting condensates near the critical concentration.

In some cases, all the proteins in the box become part of the condensate by the end of the simulations, while in other cases, there was coexistence between the dilute and condensed phases. For all the systems, the cluster size increased with concentration, mainly due to more material available to condense while the density of the dense phase remained the same (**Figure S8**). Condensates of FUS LCD in simulations with the Tesei *et al*. 2021^29^ and Dignon *et al*. 2018 models^23^ were larger and denser than the clusters in our model (**Figure S8**). The smaller condensate size with COCOMO is a result of a fraction of proteins remaining in the dilute phase, whereas the higher density in the other models may suggest overpacking of the chains during condensate formation.

We further tested our model by constructing phase diagrams as a function of temperature. Starting from a box with FUS LCD at 0.26 mM, we simulated the system at temperatures ranging from 260 to 310 K. A temperature-concentration phase diagram was constructed based on the densities in the dilute and condensed phases (**Figure 4**). We obtained very good agreement with the experimental coexistence densities reported for FUS LCD^57^. On the other hand, results for Ddx4 indicate that COCOMO may result in overpacking in the dense phase with respect to the experimental values (**Figure 4**)^58^. We note that, Regy *et al*.^28^ were able to reproduce experimental densities for this protein, and it will require further investigation to what extent correct packing can be reproduced accurately with a residue-based CG model

**Figure 4.**
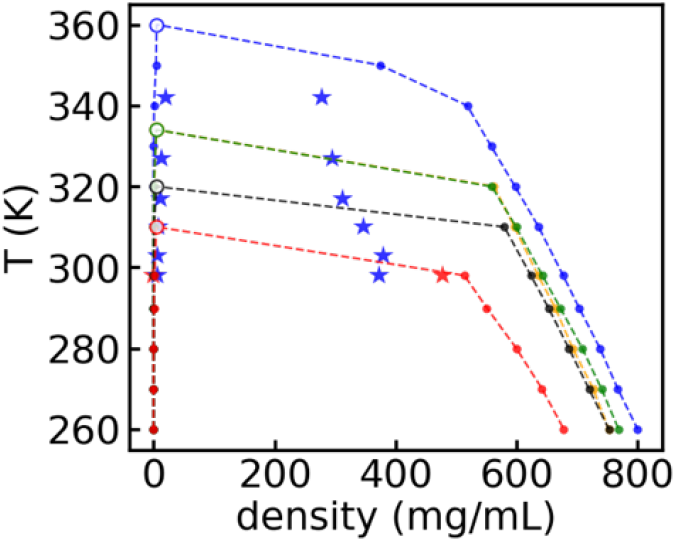
Temperature phase diagram. Simulation densities at different temperature (T) for a system starting at 4.6 mg/mL FUS LCD (red), hTau40-k18 (black), LAF-1^RGG^ (green), A1 LCD (orange), and at 5 mg/mL for Ddx4 (blue). Open circles indicate the lowest temperature where no PS was observed in the set, corresponding to *T*_*c*_. Note that LAF-1^RGG^ (green) and A1 LCD (orange) traces are almost completely overlapped. Data shown are the averaged values over the last 4 µs of the trajectory. Experimental coexistence densities for FUS LCD^57^ and Ddx4^58^ are shown using star symbols.

Recently, Tejedor *et al*.^59^ described a positive correlation between the critical temperature, *T*_*c*_, above which no LLPS is observed with the experimental saturation concentration for various phase-separating proteins, using the model developed by Dignon *et al*. (used in this work for comparison as well) with an additional reparameterization to include cation-π interactions^31^. With COCOMO, we also find such a correlation among the proteins used in the homotypic PS studies (**Figure 4**). The experimental protein saturation threshold reported are 0.013, 0.024, 0.050, and 0.24 mM for A1 LCD, LAF-1^RGG^, hTau40-k18, and FUS LCD, respectively^53-56^. The order of the experimental concentration thresholds (FUS LCD > hTau40-k18 > LAF-1^RGG^ > A1 LCD) matched the order of critical temperatures as evident from the phase diagrams (**Figure 4**). We note that the phase diagrams and *T*_*c*_ values were very similar between LAF-1^RGG^ and A1 LCD. This may be expected since the experimental saturation concentration values of these two proteins are the lowest and close to each other.

### Protein heterotypic phase separation

We further evaluated COCOMO with systems containing more than one protein as those systems can also lead to PS^53, 60, 61^. We focused on heterotypic protein PS of FUS LCD at increasing amounts of [RGRGG]_5_ and FUS LCD^RGG3^ peptides because these proteins have been well studied both experimentally and computationally^17, 28, 45, 53, 60^. Details about the simulated systems for heterotypic PS can be found in **Table S7**. Results shown in **Figure 5** indicate that COCOMO reproduces concentration-dependent PS upon addition of the peptides. As for homotypic PS, the models from Tesei *et al*. 2021^29^ and Dignon *et al*. 2018^23^ also resulted in condensation and are independent of the FUS LCD:peptide ratio. No condensation was observed with the models of Regy *et al*. 2021^28^ and Dannenhoffer *et al*. 2021^27^ at any concentration (**Figure 5** and **Figure S9**). In all cases where condensates were observed, an increase in peptide concentration was accompanied by faster cluster growth and larger final clusters (**Figure S10** and **Figure S11**).

**Figure 5.**
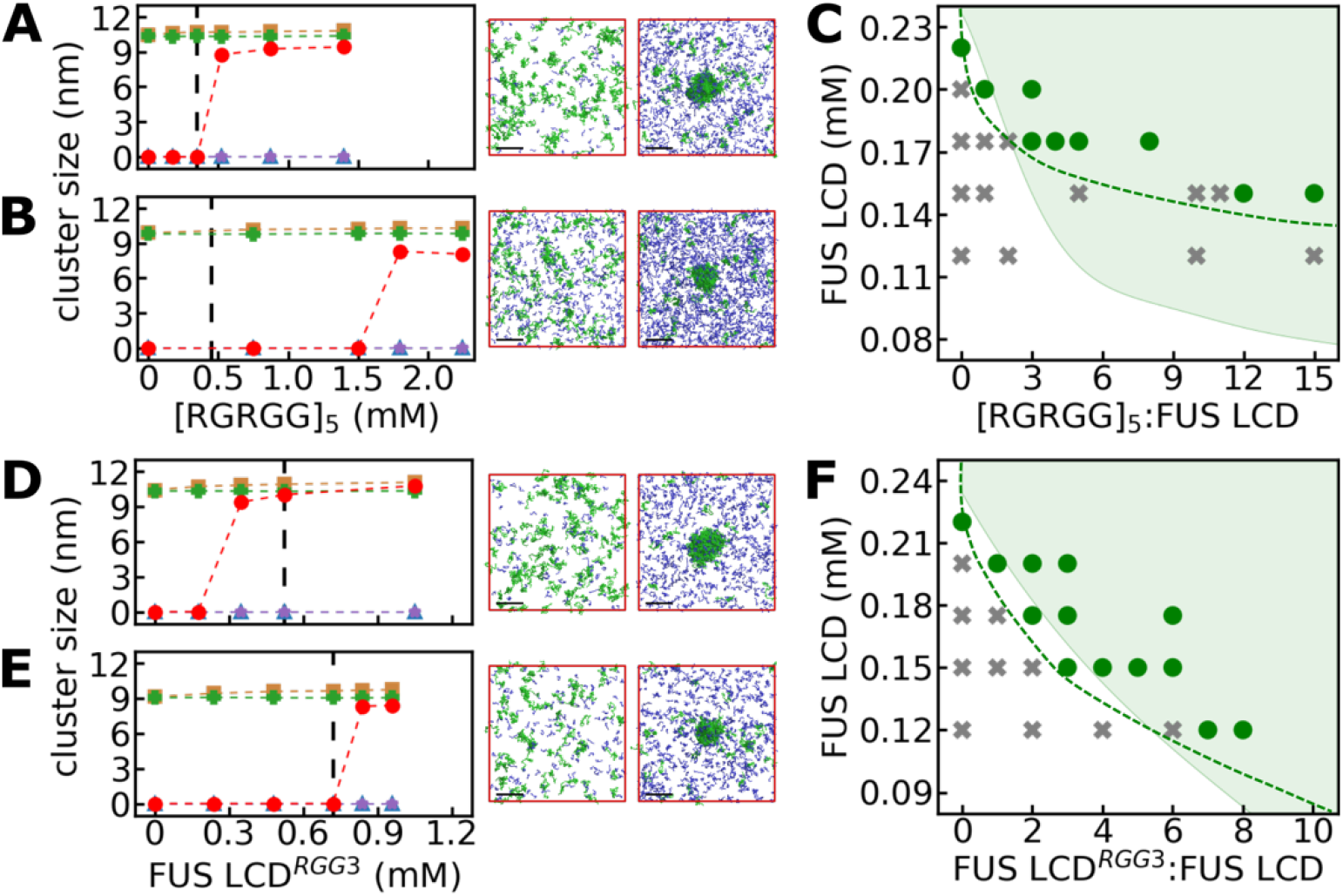
Protein heterotypic phase separation. Results are shown for FUS LCD 0.175 mM (A) and FUS LCD 0.150 mM (B) at increasing concentrations of [RGRGG]_5_ peptide, and FUS LCD 0.175 mM (C), and FUS LCD 0.120 mM (D) at increasing concentrations of FUS LCD^RGG3^ peptide. Results obtained with COCOMO (red circle) are compared with other models: Regy *et al*. 2021^28^ (blue triangle), Tesei *et al*. 2021^29^ (tan square), Dignon *et al*. 2018^23^ (green filled plus), Dannenhoffer *et al*. 2021^27^ (purple star). Cluster sizes were averaged over the last 4 µs of the simulations. The experimental LLPS concentration threshold for the addition of the peptide is shown as dashed line ^53^. Final frames of the simulations with COCOMO are shown for the lowest (left) and highest (right) concentrations added of peptide in each case. FUS LCD and peptide chains were colored in green and blue, respectively. The size bar represents 20 nm. A two-dimensional concentration-dependent phase diagram for FUS LCD as a function of different concentration ratios of [RGRGG]_5_ (C) or FUS LCD^RGG3^ (F) is shown based on COCOMO simulations. Markers show the presence (green circles) or absence (gray crosses) of condensates during simulation. The shaded region is the experimental PS regime estimated from experimental phase diagrams^53^. The dashed line is an aid to the eye for the PS regime boundary in our model.

Multicomponent phase diagrams as a function of concentration also show overall good agreement with experimental data (**Figure 5C** and **Figure 5F**). Qualitatively, the main features of the experimental phase diagrams are reproduced well. Quantitative agreement is also very good for the FUS LCD:FUS LCD^RGG3^ system (**Figure 5F**), but for the FUS LCD: [RGRGG]_5_ system, the model predicts a shift to larger minimum FUS LCD concentrations compared to the experimental data (**Figure 5C**, shaded area *vs* dashed line). Finally, the morphology of the coacervates in the simulations is also generally in agreement with experiments, as they are composed by both FUS LCD and the peptide (**Figure 5** and **Figure S10**). We found that condensates are enriched in FUS LCD over the peptide, by about tenfold, but there is no experimental data to validate the model prediction. Otherwise, our results reproduce the experimental observation that peptides enhance PS of FUS LCD^53^.

### Protein - RNA phase separation

Finally, we turn to protein-RNA condensation. Experimental evidence has shown that RNA can modulate the stability^45, 62, 63^ and kinetic properties^19, 54^ of protein condensates and protein-RNA condensation is receiving increasing attention. Therefore, another goal of our model was to describe PS in systems including proteins and nucleic acids. COCOMO was successful in capturing phase behavior of different protein-RNA mixtures for which LLPS has been described experimentally (**Figure 6** and **Table S9**). Condensates were composed of both proteins and RNA and observed in all systems that we studied. For comparison, we simulated the same systems using the model by Regy *et al*. 2020^33^. Using this model, no PS was found except for polyAde-500 – (RGRGG)_5_, where transient cluster formation was observed without clear condensation into larger clusters (**Figure 6, Figure S12**, and **Figure S13**).

**Figure 6.**
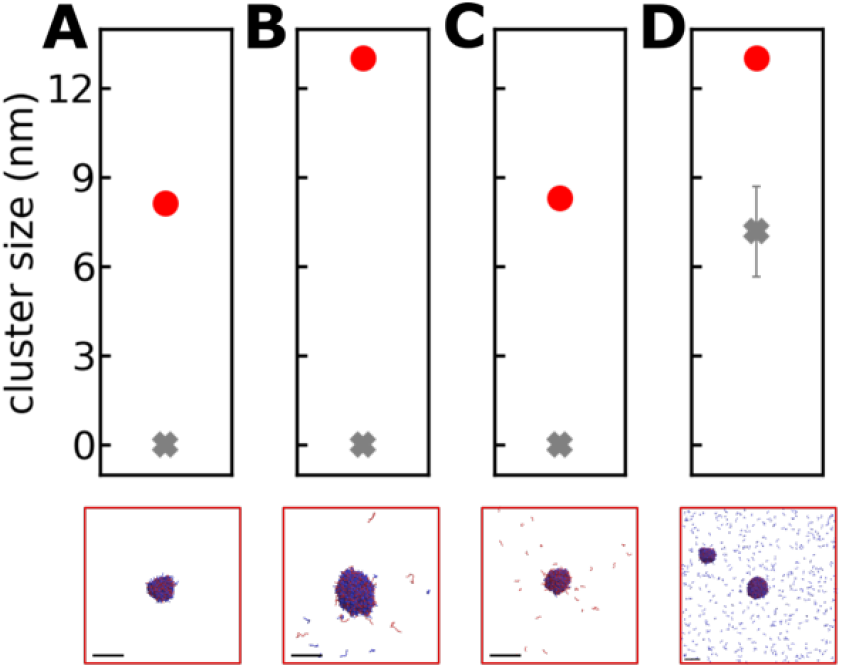
RNA – protein phase separation. Results are shown for using polyAde-21 – (RRLR)_6_-SSSGSS (A), polyUra-40 – FUS LCD^RGG3^ (B), polyUra-10 – polyArg-50 (C), and polyAde-500 – (RGRGG)_5_ (D), simulated at concentrations where PS was observed experimentally^45, 53, 64, 65^ (see details in **Table S9**). Results with COCOMO (red circle) are compared with simulations using the Regy *et al*. 2020^33^ (gray cross) model. Cluster sizes were averaged over the last 4 µs of the simulation. Final frames of simulations using COCOMO are shown on lower panels for each system. RNA and protein chains are colored in red and blue, respectively, and the size bar represents 20 nm.

We note that the choice of the value for the *ε*_*R*/*K* − *nucleic*_ parameter affects the agreement with the experimental data^45, 53, 64, 65^. Previous studies showed contrasting views on this term, Das *et al*.^31^ argued that despite augmenting *ε*_*R*/*K* − *nucleic*_ parameter, their model failed to capture the experimental PS propensity, while other authors^59^ demonstrated that increased cation-π interactions can reproduce the experimental trend of their selected set of proteins. Here, we find that with this interaction added explicitly we can describe PS between proteins and RNA in agreement with experiments.

## CONCLUSIONS

In this work, we present a simple coarse-grained model that can balance polymer properties of disordered proteins and RNA with their propensity to phase separate. Our model differs from the other four used for comparison^23, 27-29^ in various key aspects, the presence of an angle term to account for protein chain stiffness, the introduction of a term for cation-π interactions, and the implicit description of solvation effects. As found in previous studies^26^, we find that enhanced cation-π interaction is an important aspect of the model.

Our model reproduces experimental *R*_*g*_ values for peptides well and captures the dependency of RNA polymer properties on salt concentration. At the same time, the model can describe homotypic and heterotypic protein PS as well as PS involving protein-RNA mixtures in agreement with various *in vitro* experimental systems. Other models can also accurately reproduce *R*_*g*_ values for different peptide sequences, and some of those models can also describe PS. However, in those models, concentration-dependent PS involves significantly lower temperatures or higher concentrations than what is reported in the experiments. The key strength of the model here is that experimental behavior is reproduced at the level of single polymers as well as for condensation at the same temperature and in a concentration-dependent manner that matches experiments. Therefore, the model is more predictive with respect to when and under what conditions PS would be expected for systems for which experimental data is not available.

Even though the COCOMO CG model presented here performs well when compared to experimental observations, there are also significant limitations. The implicit description of salt effects based on the Debye-Hückel formalism is only approximate and does not capture changes in free energy of the ions themselves as they condense along with the biopolymers. In addition, the residue-level approximation neglects any shape-dependent packing interactions during condensation, and, more generally, anisotropic or directional interactions, for example due to aromatic stacking or hydrogen bonding. The next steps to further improve the model would be to convert condensate models generated by the COCOMO to higher-resolution representations and test their physical viability via simulation. For example, atomistic simulations of reconstructed condensates may inform how to improve the CG model without increasing the computational complexity.

A physically realistic and computationally efficient CG model that is predictive and can reach the time and spatial scales on which biomolecular condensation is observed experimentally opens up a wide range of applications. Despite much experimental information obtained so far about biomolecular condensates, many questions about how condensation depends on molecular compositions remain unclear. The model here allows such questions to be explored, not just for homo- and heterotypic peptide systems, but also for peptide-RNA mixtures. We expect that there will be many applications where such simulations can interpret experimental observations and suggest new hypotheses to be tested experimentally.

## ACKNOWLEDGEMENTS

Funding was provided by the National Science Foundation grants MCB 1817307 and by the National Institute of Health (NIGMS) grant R35 GM126948. We used computational resources at the Institute for Cyber-Enabled Research/High Performance Computing Cluster (ICER/HPCC) at Michigan State University.

## REFERENCES

(1) Shin, Y.; Brangwynne, C. P. Liquid phase condensation in cell physiology and disease. Science 2017, 357, eaaf4382.

(2) Pederson, T. The nucleolus. Cold Spring Harbor Perspect. Biol. 2011, 3, a000638.

(3) Mao, Y. S.; Zhang, B.; Spector, D. L. Biogenesis and function of nuclear bodies. Trends Genet. 2011, 27, 295–306.

(4) Gall, J. G. Cajal bodies: the first 100 years. Annu. Rev. Cell Dev. Biol. 2000, 16, 273–300.

(5) Gall, J. G. The centennial of the Cajal body. Nat. Rev. Mol. Cell Biol. 2003, 4, 975–980.

(6) Lamond, A. I.; Spector, D. L. Nuclear speckles: A model for nuclear organelles. Nat. Rev. ol. Cell Biol. 2003, 4, 605–612.

(7) Galganski, L.; Urbanek, M. O.; Krzyzosiak, W. J. Nuclear speckles: molecular organization, biological function and role in disease. Nucleic Acids Res. 2017, 45, 10350–10368.

(8) Buchan, J. R.; Parker, R. Eukaryotic stress granules: the ins and outs of translation. Mol. Cell 2009, 36, 932–941.

(9) Protter, D. S. W.; Parker, R. Principles and Properties of Stress Granules. Trends Cell Biol. 2016, 26, 668–679.

(10) Luo, Y.; Na, Z.; Slavoff, S. A. P-Bodies: Composition, Properties, and Functions. Biochemistry 2018, 57, 2424–2431.

(11) Voronina, E.; Seydoux, G.; Sassone-Corsi, P.; Nagamori, I. RNA Granules in Germ Cells. Cold Spring Harbor Perspect. Biol. 2011, 3, a002774.

(12) Lyon, A. S.; Peeples, W. B.; Rosen, M. K. A framework for understanding the functions of biomolecular condensates across scales. Nat. Rev. Mol. Cell Biol. 2021, 22, 215–235.

(13) Banani, S. F.; Lee, H. O.; Hyman, A. A.; Rosen, M. K. Biomolecular condensates: organizers of cellular biochemistry. Nat. Rev. Mol. Cell Biol. 2017, 18, 285–298.

(14) Mitrea, D. M.; Chandra, B.; Ferrolino, M. C.; Gibbs, E. B.; Tolbert, M.; White, M. R.; Kriwacki, R. W. Methods for Physical Characterization of Phase-Separated Bodies and Membrane-less Organelles. J. Mol. Biol. 2018, 430, 4773–4805.

(15) Brangwynne, C. P.; Tompa, P.; Pappu, R. V. Polymer physics of intracellular phase transitions. Nat. Phys. 2015, 11, 899–904.

(16) Posey, A. E.; Holehouse, A. S.; Pappu, R. V. Phase Separation of Intrinsically Disordered Proteins. Methods Enzymol. 2018, 611, 1–30.

(17) Zheng, W.; Dignon, G. L.; Jovic, N.; Xu, X.; Regy, R. M.; Fawzi, N. L.; Kim, Y. C.; Best, R. B.; Mittal, J. Molecular Details of Protein Condensates Probed by Microsecond Long Atomistic Simulations. J. Phys. Chem. B 2020, 124, 11671–11679.

(18) Paloni, M.; Bailly, R.; Ciandrini, L.; Barducci, A. Unraveling Molecular Interactions in Liquid-Liquid Phase Separation of Disordered Proteins by Atomistic Simulations. J. Phys. Chem. B 2020, 124, 9009–9016.

(19) Wei, M. T.; Elbaum-Garfinkle, S.; Holehouse, A. S.; Chen, C. C.; Feric, M.; Arnold, C. B.; Priestley, R. D.; Pappu, R. V.; Brangwynne, C. P. Phase behaviour of disordered proteins underlying low density and high permeability of liquid organelles. Nat. Chem. 2017, 9, 1118–1125.

(20) Feig, M.; Chocholousova, J.; Tanizaki, S. Extending the Horizon: Towards the Efficient Modeling of Large Biomolecular Complexes in Atomic Detail. Theor. Chem. Acc. 2006, 116, 194–205.

(21) Predeus, A. V.; Gul, S.; Gopal, S. M.; Feig, M. Conformational Sampling of Peptides in the Presence of Protein Crowders from AA/CG-Multiscale Simulations. J. Phys. Chem. B 2012, 116, 8610–8620.

(22) Kar, P.; Feig, M. Recent Advances in Transferable Coarse-Grained Modeling of Proteins. Adv. Prot. Chem. Struct. Biol. 2014, 96, 143–180.

(23) Dignon, G. L.; Zheng, W.; Kim, Y. C.; Best, R. B.; Mittal, J. Sequence determinants of protein phase behavior from a coarse-grained model. Plos Comp. Biol. 2018, 14, e1005941.

(24) Mani, E.; Lechner, W.; Kegel, W. K.; Bolhuis, P. G. Equilibrium and Non-Equilibrium Cluster Phases in Colloids with Competing Interactions. Soft Matter 2014, 10, 4479–4486.

(25) Dutagaci, B.; Nawrocki, G.; Goodluck, J.; Ashkarran, A. A.; Hoogstraten, C. G.; Lapidus, L. J.; Feig, M. Charge-driven condensation of RNA and proteins suggests broad role of phase separation in cytoplasmic environments. eLife 2021, 10, e64004.

(26) Joseph, J. A.; Espinosa, J. R.; Sanchez-Burgos, I.; Garaizar, A.; Frenkel, D.; Collepardo-Guevara, R. Thermodynamics and kinetics of phase separation of protein-RNA mixtures by a minimal model. Biophys. J. 2021, 120, 1219–1230.

(27) Dannenhoffer-Lafage, T.; Best, R. B. A Data-Driven Hydrophobicity Scale for Predicting Liquid-Liquid Phase Separation of Proteins. J. Phys. Chem. B 2021, 125, 4046–4056.

(28) Regy, R. M.; Thompson, J.; Kim, Y. C.; Mittal, J. Improved coarse-grained model for studying sequence dependent phase separation of disordered proteins. Protein Sci. 2021, 30, 1371–1379.

(29) Tesei, G.; Schulze, T. K.; Crehuet, R.; Lindorff-Larsen, K. Accurate model of liquid-liquid phase behavior of intrinsically disordered proteins from optimization of single-chain properties. Proc. Natl. Acad. Sci. U.S.A. 2021, 118, e2111696118.

(30) Latham, A. P.; Zhang, B. Maximum Entropy Optimized Force Field for Intrinsically Disordered Proteins. J. Chem. Theory Comput. 2020, 16, 773–781.

(31) Das, S.; Lin, Y. H.; Vernon, R. M.; Forman-Kay, J. D.; Chan, H. S. Comparative roles of charge, pi, and hydrophobic interactions in sequence-dependent phase separation of intrinsically disordered proteins. Proc. Natl. Acad. Sci. U.S.A. 2020, 117, 28795–28805.

(32) Joseph, J. A.; Reinhardt, A.; Aguirre, A.; Chew, P. Y.; Russell, K. O.; Espinosa, J. R.; Garaizar, A.; Collepardo-Guevara, R. Physics-driven coarse-grained model for biomolecular phase separation with near-quantitative accuracy. Nat. Comput. Sci. 2021, 1, 732–743.

(33) Regy, R. M.; Dignon, G. L.; Zheng, W. W.; Kim, Y. C.; Mittal, J. Sequence dependent phase separation of protein-polynucleotide mixtures elucidated using molecular simulations. Nucleic Acids Res. 2020, 48, 12593–12603.

(34) Wu, H.; Wolynes, P. G.; Papoian, G. A. AWSEM-IDP: A Coarse-Grained Force Field for Intrinsically Disordered Proteins. J. Phys. Chem. B 2018, 122, 11115–11125.

(35) Holmstrom, E. D.; Liu, Z.; Nettels, D.; Best, R. B.; Schuler, B. Disordered RNA chaperones can enhance nucleic acid folding via local charge screening. Nat. Commun. 2019, 10, 2453.

(36) Marrink, S. J.; Risselada, H. J.; Yefimov, S.; Tieleman, D. P.; de Vries, A. H. The MARTINI force field: coarse grained model for biomolecular simulations. J. Phys. Chem. B 2007, 111, 7812–7824.

(37) Ashbaugh, H. S.; Hatch, H. W. Natively unfolded protein stability as a coil-to-globule transition in charge/hydropathy space. J. Am. Chem. Soc. 2008, 130, 9536–9542.

(38) Farahi, N.; Lazar, T.; Wodak, S. J.; Tompa, P.; Pancsa, R. Integration of Data from Liquid-Liquid Phase Separation Databases Highlights Concentration and Dosage Sensitivity of LLPS Drivers. Int. J. Mol. Sci. 2021, 22.

(39) Ghobadi, A. F.; Jayaraman, A. Effect of backbone chemistry on hybridization thermodynamics of oligonucleic acids: a coarse-grained molecular dynamics simulation study. Soft Matter 2016, 12, 2276–2287.

(40) Eastman, P.; Swails, J.; Chodera, J. D.; McGibbon, R. T.; Zhao, Y.; Beauchamp, K. A.; Wang, L. P.; Simmonett, A. C.; Harrigan, M. P.; Stern, C. D.; et al. OpenMM 7: Rapid development of high performance algorithms for molecular dynamics. Plos Comp. Biol. 2017, 13, e1005659.

(41) Feig, M.; Karanicolas, J.; Brooks, C. L. MMTSB Tool Set: Enhanced Sampling and Multiscale Modeling Methods for Applications in Structural Biology. J. Mol. Graph. Modell. 2004, 22, 377–395.

(42) Brooks, B. R.; Bruccoleri, R. E.; Olafson, B. D.; States, D. J.; Swaminathan, S.; Karplus, M. Charmm - a Program for Macromolecular Energy, Minimization, and Dynamics Calculations. J. Comput. Chem. 1983, 4, 187–217.

(43) McGibbon, R. T.; Beauchamp, K. A.; Harrigan, M. P.; Klein, C.; Swails, J. M.; Hernandez, C. X.; Schwantes, C. R.; Wang, L. P.; Lane, T. J.; Pande, V. S. MDTraj: A Modern Open Library for the Analysis of Molecular Dynamics Trajectories. Biophys. J. 2015, 109, 1528–1532.

(44) Beauchamp, K. A.; Bowman, G. R.; Lane, T. J.; Maibaum, L.; Haque, I. S.; Pande, V. S. MSMBuilder2: Modeling Conformational Dynamics at the Picosecond to Millisecond Scale. J. Chem. Theory Comput. 2011, 7, 3412–3419.

(45) Alshareedah, I.; Kaur, T.; Ngo, J.; Seppala, H.; Kounatse, L. D.; Wang, W.; Moosa, M. M.; Banerjee, P. R. Interplay between Short-Range Attraction and Long-Range Repulsion Controls Reentrant Liquid Condensation of Ribonucleoprotein-RNA Complexes. J. Am. Chem. Soc. 2019, 141, 14593–14602.

(46) Fisher, R. S.; Elbaum-Garfinkle, S. Tunable multiphase dynamics of arginine and lysine liquid condensates. Nat. Commun. 2020, 11, 4628.

(47) Dignon, G. L.; Best, R. B.; Mittal, J. Biomolecular Phase Separation: From Molecular Driving Forces to Macroscopic Properties. Annu. Rev. Phys. Chem. 2020, 71, 53–75.

(48) Paloni, M.; Bussi, G.; Barducci, A. Arginine multivalency stabilizes protein/RNA condensates. Protein Sci. 2021, 30, 1418–1426.

(49) Ukmar-Godec, T.; Hutten, S.; Grieshop, M. P.; Rezaei-Ghaleh, N.; Cima-Omori, M. S.; Biernat, J.; Mandelkow, E.; Soding, J.; Dormann, D.; Zweckstetter, M. Lysine/RNA-interactions drive and regulate biomolecular condensation. Nat. Commun. 2019, 10, 2909.

(50) Plumridge, A.; Andresen, K.; Pollack, L. Visualizing Disordered Single-Stranded RNA: Connecting Sequence, Structure, and Electrostatics. J. Am. Chem. Soc. 2020, 142, 109–119.

(51) Borgia, A.; Borgia, M. B.; Bugge, K.; Kissling, V. M.; Heidarsson, P. O.; Fernandes, C. B.; Sottini, A.; Soranno, A.; Buholzer, K. J.; Nettels, D.; et al. Extreme disorder in an ultrahigh-affinity protein complex. Nature 2018, 555, 61–66.

(52) Chen, H.; Meisburger, S. P.; Pabit, S. A.; Sutton, J. L.; Webb, W. W.; Pollack, L. Ionic strength-dependent persistence lengths of single-stranded RNA and DNA. Proc. Natl. Acad. Sci. U.S.A. 2012, 109, 799–804.

(53) Kaur, T.; Raju, M.; Alshareedah, I.; Davis, R. B.; Potoyan, D. A.; Banerjee, P. R. Sequence-encoded and composition-dependent protein-RNA interactions control multiphasic condensate morphologies. Nat. Commun. 2021, 12, 872.

(54) Elbaum-Garfinkle, S.; Kim, Y.; Szczepaniak, K.; Chen, C. C. H.; Eckmann, C. R.; Myong, S.; Brangwynne, C. P. The disordered P granule protein LAF-1 drives phase separation into droplets with tunable viscosity and dynamics. Proc. Natl. Acad. Sci. U.S.A. 2015, 112, 7189–7194.

(55) Ambadipudi, S.; Biernat, J.; Riedel, D.; Mandelkow, E.; Zweckstetter, M. Liquid-liquid phase separation of the microtubule-binding repeats of the Alzheimer-related protein Tau. Nat. Commun. 2017, 8, 275.

(56) Bremer, A.; Farag, M.; Borcherds, W. M.; Peran, I.; Martin, E. W.; Pappu, R. V.; Tanja, M. Deciphering how naturally occurring sequence features impact the phase behaviours of disordered prion-like domains. Nat. Chem. 2022, 14, 196--207.

(57) Murthy, A. C.; Dignon, G. L.; Kan, Y.; Zerze, G. H.; Parekh, S. H.; Mittal, J.; Fawzi, N. L. Molecular interactions underlying liquid-liquid phase separation of the FUS low-complexity domain. Nat. Struct. Mol. Biol. 2019, 26, 637–648.

(58) Brady, J. P.; Farber, P. J.; Sekhar, A.; Lin, Y. H.; Huang, R.; Bah, A.; Nott, T. J.; Chan, H. S.; Baldwin, A. J.; Forman-Kay, J. D.; et al. Structural and hydrodynamic properties of an intrinsically disordered region of a germ cell-specific protein on phase separation. Proc. Natl. Acad. Sci. U.S.A. 2017, 114, E8194–E8203.

(59) Tejedor, A. R.; Garaizar, A.; Ramirez, J.; Espinosa, J. R. RNA modulation of transport properties and stability in phase-separated condensates. Biophys. J. 2021, 120, 5169–5186.

(60) Boeynaems, S.; Bogaert, E.; Kovacs, D.; Konijnenberg, A.; Timmerman, E.; Volkov, A.; Guharoy, M.; De Decker, M.; Jaspers, T.; Ryan, V. H.; et al. Phase Separation of C9orf72 Dipeptide Repeats Perturbs Stress Granule Dynamics. Mol. Cell 2017, 65, 1044–1055 e1045.

(61) Sanders, D. W.; Kedersha, N.; Lee, D. S. W.; Strom, A. R.; Drake, V.; Riback, J. A.; Bracha, D.; Eeftens, J. M.; Iwanicki, A.; Wang, A.; et al. Competing Protein-RNA Interaction Networks Control Multiphase Intracellular Organization. Cell 2020, 181, 306–324 e328.

(62) Schwartz, J. C.; Wang, X.; Podell, E. R.; Cech, T. R. RNA seeds higher-order assembly of FUS protein. Cell Rep. 2013, 5, 918–925.

(63) Banerjee, P. R.; Milin, A. N.; Moosa, M. M.; Onuchic, P. L.; Deniz, A. A. Reentrant Phase Transition Drives Dynamic Substructure Formation in Ribonucleoprotein Droplets. Angew. Chemie Int. Ed. 2017, 56, 11354–11359.

(64) Bai, Q.; Zhang, Q.; Jing, H.; Chen, J.; Liang, D. Liquid-Liquid Phase Separation of Peptide/Oligonucleotide Complexes in Crowded Macromolecular Media. J. Phys. Chem. B 2021, 125, 49–57.

(65) Fisher, R. S.; Elbaum-Garfinkle, S. Tunable multiphase dynamics of arginine and lysine liquid condensates. Nat. Commun. 2020, 11, 4628.

