## Supplementary figures and tables for "Modeling concentration-dependent phase separation processes involving peptides and RNA via residue-based coarse-graining"

for

**Tables S1-S9**
**Figures S1-S13**
**Supplementary References**

#### Table S1. Residue-specific parameters used in COCOMO.

| Residue | Mass(amu) | $r_i$ (nm)† | charge | $A_i$‡ | $A0_i$ |
|---|---|---|---|---|---|
| Ala | 71.08 | 0.2845 | 0 | 0 | 0 |
| Arg | 157.20 | 0.3567 | 1 | 0.866 | 0.05 |
| Asn | 114.10 | 0.3150 | 0 | 0 | 0.05 |
| Asp | 114.08 | 0.3114 | -1 | -0.866 | 0.05 |
| Cys | 103.14 | 0.3024 | 0 | 0 | 0.05 |
| Gln | 128.13 | 0.3311 | 0 | 0 | 0.05 |
| Glu | 128.11 | 0.3279 | -1 | -0.866 | 0.05 |
| Gly | 57.05 | 0.2617 | 0 | 0 | 0 |
| His | 137.14 | 0.3338 | 0 | 0 | 0.05 |
| Ile | 113.16 | 0.3360 | 0 | 0 | 0 |
| Leu | 113.16 | 0.3363 | 0 | 0 | 0 |
| Lys | 129.18 | 0.3439 | 1 | 0.866 | 0.05 |
| Met | 131.19 | 0.3381 | 0 | 0 | 0 |
| Phe | 147.18 | 0.3556 | 0 | 0 | 0 |
| Pro | 98.13 | 0.3187 | 0 | 0 | 0 |
| Ser | 87.08 | 0.2927 | 0 | 0 | 0.05 |
| Thr | 101.11 | 0.3108 | 0 | 0 | 0.05 |
| Trp | 186.21 | 0.3754 | 0 | 0 | 0 |
| Tyr | 71.08 | 0.2845 | 0 | 0 | 0 |
| Val | 157.20 | 0.3567 | 1 | 0 | 0 |
| Ade | 315.697 | 0.4220 | -1 | -0.866 | 0.05 |
| Cyt | 305.200 | 0.4110 | -1 | -0.866 | 0.05 |
| Gua | 345.200 | 0.4255 | -1 | -0.866 | 0.05 |
| Ura | 305.162 | 0.4090 | -1 | -0.866 | 0.05 |

† $r_i$, radius of a sphere with equivalent volume of a given residue.

‡ $A_i = \text{sign}(q_i)\sqrt{0.75|q_i|}$, where $q_i$ is the residue net charge.

#### Table S2. Sequences of disordered or unfolded proteins in the parametrization set.

| | |
|---|---|
| ACTR | GTQNRPLLRN SLDDLVGPPS NLEGQSDERA LLDQLHTLLS NTDATGLEEI DRALGIPELV NQGQALEPKQ D |
| ak16 | YGCAKAAAAK ACAAKA |
| ak27 | AAKAAAAKAA AAKAAAAKAA AAKAAGY |
| ak32 | AAKAAAAKAA AAKAAAAKAA AAKAAAAKAA GY |
| ak37 | AAKAAAAKAA AAKAAAAKAA AAKAAAAKAA AAKAAGY |
| AN16 | MHHHHHHPGA PAQTPSSQYG APAQTPSSQY GAPAQTPSSQ YGAPAQTPSS QYGAPAQTPS SQYGAPAQTP SSQYGAPAQT PSSQYGAPAQ TPSSQYGAPA QTPSSQYGAP AQTPSSQYGA PAQTPSSQYG APAQTPSSQY GAPAQTPSSQ YGAPAQTPSS QYGAPAQTPS SQYGAPAQTP SSQYV |
| angiotensin | DRVYIHPF |
| α-synuclein | MDVFMKGLSK AKEGVVAAAE KTKQGVAEAA GKTKEGVLYV GSKTKEGVVH GVATVAEKTK EQVTNVGGAV VTGVTAVAQK TVEGAGSIAA ATGFVKKDQL GKNEEGAPQE GILEDMPVDP DNEAYEMPSE EGYQDYEPEA |
| CspTm | GPGMRGKVKW FDSKKGYGFI TKDEGGDVFV HWSAIEMEGF KTLKEGQVVE FEIQEGKKGG QAAHVKV |
| cytC-nter | MIFFMVMPIM IGGFGNWLVP LMIGAPDMAF PRMNNSFWL |
| drK-SH3 | MEAIAKHDFS ATADDELSFR KTQILKILNM EDDSNWYRAE LDGKEGLIPS NYIEMKNHD |
| Erm | MDGFYDQQVP FMVPGKSRSE ECRGRPVIDR KRKFLDTDLA HDSEELFQDL SQLQEAWLAE AQVPDDEQFV PDFQSDNLVL HAPPPTKIKR ELHSPSSELS SCSHEQALGA NYGEKCLYNY CA |
| Fhua | ESAWGPAATI AARQSATGTK TDTPIQKVPQ SISVVTAEEM ALHQPKSVKE ALSYTPGVSV GTRGASNTYD HLIIRGFAAE GQSQNNYLNG LKLQGNFYND AVIDPYMLER AEIMRGPVSV LYGKSSPGGL LNMVSKRPTT EPL |
| hCyp | GPMCNPTVFF DIAVDGEPLG RVSFELFADK VPKTAENFRA LSTGEKGFGY KGSSFHRIIP GFMSQGGDFT RHNGTGGKSI YGEKFEDENF ILKHTGPGIL SMANAGPNTN GSQFFISTAK TEFLDGKHVV FGKVKEGMNI VEAMERFGSR NGKTSKKITI ADSGQLC |
| His5 | DSHAKRHHGY KRKFHEKHHS HRGY |
| hNHE1cdt | MVPAHKLDSP TMSRARIGSD PLAYEPKEDL PVITIDPASP QSPESVDLVN EELKGKVLGL SRDPAKVAEE DEDDDGGIMM RSKETSSPGT DDVFTPAPSD SPSSQRIQRC LSDPGPHPEP GEGEPFFPKG Q |
| hTau23-k10 | MQTAPVPMPD LKNVKSKIGS TENLKHQPGG GKVQIVYKPV DLSKVTSKCG SLGNIHHKPG GGQVEVKSEK LDFKDRVQSK IGSLDNITHV PGGGNKKIET HKLTFRENAK AKTDHGAEIV YKSPVVSGDT SPRHLSNVSS TGSIDMVDSP QLATLADEVS ASLAKQGL |
| hTau23-k17 | MSSPGSPGTP GSRSRTPSLP TPPTREPKKV AVVRTPPKSP SSAKSRLQTA PVPMPDLKNV KSKIGSTENL KHQPGGGKVQ IVYKPVDLSK VTSKCGSLGN IHHKPGGGQV EVKSEKLDFK DRVQSKIGSL DNITHVPGGG NKKIE |
| hTau23-k19 | MQTAPVPMPD LKNVKSKIGS TENLKHQPGG GKVQIVYKPV DLSKVTSKCG SLGNIHHKPG GGQVEVKSEK LDFKDRVQSK IGSLDNITHV PGGGNKKIE |
| hTau23-k25 | MAEPRQEFEV MEDHAGTYGL GDRKDQGGYT MHQDQEGDTD AGLKAEEAGI GDTPSLEDEA AGHVTQARMV SKSKDGTGSD DKKAKGADGK TKIATPRGAA PPGQKGQANA TRIPAKTPPA PKTPPSSGEP PKSGDRSGYS SPGSPGTPGS RSRTPSLPTP PTREPKKVAV VRTPPKSPSS AKSRL |
| hTau23-k27 | MSSPGSPGTP GSRSRTPSLP TPPTREPKKV AVVRTPPKSP SSAKSRLQTA PVPMPDLKNV KSKIGSTENL KHQPGGGKVQ IVYKPVDLSK VTSKCGSLGN IHHKPGGGQV EVKSEKLDFK DRVQSKIGSL DNITHVPGGG NKKIETHKLT FRENAKAKTD HGAEIVY |
| hTau40-k16 | MSSPGSPGTP GSRSRTPSLP TPPTREPKKV AVVRTPPKSP SSAKSRLQTA PVPMPDLKNV KSKIGSTENL KHQPGGGKVQ IINKKLDLSN VQSKCGSKDN IKHVPGGGSV QIVYKPVDLS KVTSKCGSLG NIHHKPGGGQ VEVKSEKLDF KDRVQSKIGS LDNITHVPGG GNKKIE |
| hTau40-k18 | MQTAPVPMPD LKNVKSKIGS TENLKHQPGG GKVQIINKKL DLSNVQSKCG SKDNIKHVPG GGSVQIVYKP VDLSKVTSKC GSLGNIHHKP GGGQVEVKSE KLDFKDRVQS KIGSLDNITH VPGGGNKKIE |

| | |
|---|---|
| hTau40-k32 | MSSPGSPGTP GSRSRTPSLP TPPTREPKKV AVVRTPPKSP SSAKSRLQTA PVPMPDLKNV KSKIGSTENL KHQPGGGKVQ IINKKLDLSN VQSKCGSKDN IKHVPGGGSV QIVYKPVDLS KVTSKCGSLG NIHHKPGGGQ VEVKSEKLDF KDRVQSKIGS LDNITHVPGG GNKKIETHKL TFRENAKAKT DHGAEIVY |
| ibb | GCTNENANTP AARLHRFKNK GKDSTEMRRR RIEVNVELRK AKKDDQMLKR RNVSSFPDDA TSPLQENRNN QGTVNWSVDD IVKGINSSNV ENQLQATFA |
| in | GSHCFLDGID KAQEEHEKYH SNWRAMASDF NLPPVVAKEI VASCDKCQLK GEAMHGQVDC |
| n49 | GCQTSRGLFG NNNTNNINNS SSGMNNASAG LFGSKPFA |
| nls | ACETNKRKRE QISTDNEAKM QIQEEKSPKK KRKKRSSKAN KPPEFA |
| nul | GCGFKGFDTS SSSSNSAASS SFKFGVSSSS SGPSQTLTST GNFKFGDQGG FKIGVSSDSG SINPMSEGFK FSKPIGDFKF GVSSESKPEE VKKDSKNDNF KFGLSSGLSN PVFA |
| nus | GCPSASPAFG ANQTPTFGQS QGASQPNPPG FGSISSSTAL FPTGSQPAPP TFGTVSSSSQ PPVFGQQPSQ SAFGSGTTPN FA |
| p53 | MEEPQSDPSV EPPLSQETFS DLWKLLPENN VLSPLPSQAM DDLMLSPDDI EQWFTEDPGP DEAPRMPEAA PPVAPAPAAP TPAAPAPAPS WPL |
| ProT$\alpha$-C | CEEGGEEEEE EEEGDGEEED GDEDEEAESA TGKRAAEDDE DDDVDTKKQK TDEDD |
| ProT$\alpha$-N | SDAAVDTSSE ITTKDLKEKK EVVEEAENGR DAPANGNANE ENGEQEADNE VDEEC |
| Protein G | MQYKLALNGK TLKGETTTEA VDAATAEKVF KQYANDNGVD GEWAYDDATK TFAVTE |
| Protein L | MEEVTIKANL IFANGSTQTA EFKGTFEKAT SEAYAYADTL KKDNGEWTVD VADKGYTLNI KFAG |
| R15 | KLKEACKQQN FNTGIKDFDF WLSEVEALLA SEDYGKDLAS VNNLLKKHQL LEADISAHED RLKDLNSQAD SLMTSSAFDT SQVKDKRETI NGRFQRIKCM AAARRAKLNE SHRL |
| R17 | GSRLEESCEY QQFVANVEEE EAWINEKMTL VASEDYGDTL AAIQGLLKKH EAFETDFTVH KDRVNDVAAN GEDLIKKNNH HVENITAKMK GLKGKVSDLE CAAAQRKAKL DENSAFLQ |
| RNaseA | KETAAAKFER QHMDSSTSAA SSSNYCNQMM KSRNLTKDRC KPVNTFVHES LADVQAVCSQ KNVACKNGQT NCYQSYSTMS ITDCRETGSS KYPNCAYKTT QANKHIIVAC EGNPYVPVHF DASV |
| SH4 | MGSNKSKPKD ASQRRRSLEP AENVHGAGGG AFPASQTPSK PASADGHRGP SAAFAPAAAE PKLFGGFNSS DTVTSPQRAG PLAGG |
| Sic1 | MTPSTPPRSR GTRYLAQPSG NTSSSALMQG QKTPQKPSQN LVPVTPSTTK SFKNAPLLAP PNSNMGMTSP FNGLTSPQRS PFPKSSVKRT |
| sNase | ATSTKKLHKE PATLIKAIDG DTVKLMYKGQ PMTFRLLLVD TPETKHPKKG VEKYGPEASA FTKKMVENAK KIEVEFDKGQ RTDKYGRGLA YIYADGKMVN EALVRQGLAK VAYVYKPNNT HEQHLRKSEA QAKKEK |
| ubiquitin | MQIFVKTLTG KTITLEVEPS DTIENVKAKI QDKEGIPPDQ QRLIWAGKQL EDGRTLSDYN IQKESTLHLV LRLRGG |
| ul11 | MGLSFSGTRP CCCRNNVLIT DDGEVVSLTA HDFDVVDIES EEEGNFYVPP DMRGVTRAPG RQRLRSSDPP SRHTHRRTPG GACPATQFPP PMSDSEWSHP QFEK |
| yesg2 | YESGGATD |
| yesg6 | YESGGGGGGA TD |

### Table S3. Radii of gyration of proteins in the parametrization set.

| protein | Rg (nm) | [ion] (M) | pH | Method | Reference |
|---|---|---|---|---|---|
| ACTR | 2.51 | 0.20 | 7.4 | SAXS | Kjaergaard *et al.*[1] |
| ak16 | 0.98 | - | - | SAXS | Kohn *et al.*[2] |
| ak27 | 1.28 | - | - | SAXS | Kohn *et al.*[2] |
| ak32 | 1.45 | - | - | SAXS | Kohn *et al.*[2] |
| ak37 | 1.69 | - | - | SAXS | Kohn *et al.*[2] |
| AN16 | 4.44 | - | - | SAXS | Kohn *et al.*[2] |
| angiotensin | 0.79 | 0.20 | - | SAXS | Ohnishi *et al.*[3] |
| $\alpha$-synuclein | 3.30 | 0.16 | 7.4 | FRET | Nath *et al.*[4] |
| CspTm | 1.47 | 0.04 | - | FRET | Muller-Spath *et al.*[5] |
| cytC-nter | 1.84 | - | - | SAXS | Kohn *et al.*[2] |
| drK-SH3 | 2.19 | - | - | SAXS | Choy *et al.*[6] |
| erm | 3.96 | 0.20 | 8.0 | SAXS | Lens *et al.*[7] |
| fhua | 3.34 | 0.15 | 7.5 | SAXS | Riback *et al.*[8] |
| hCyp | 2.51 | 0.09 | - | FRET | Hofmann *et al.*[9] |
| His5 | 1.38 | 0.15 | 7.5 | SAXS | Cragnell *et al.*[10] |
| hNHE1cdt | 3.63 | 0.20 | - | SAXS | Kjaergaard *et al.*[1] |
| hTau23-k10 | 4.00 | 0.15 | 7.4 | SAXS | Mylonas *et al.*[11] |
| hTau23-k17 | 3.60 | 0.15 | 7.4 | SAXS | Mylonas *et al.*[11] |
| hTau23-k19 | 3.50 | 0.15 | 7.4 | SAXS | Mylonas *et al.*[11] |
| hTau23-k25 | 4.10 | 0.15 | 7.4 | SAXS | Mylonas *et al.*[11] |
| hTau23-k27 | 3.70 | 0.15 | 7.4 | SAXS | Mylonas *et al.*[11] |
| hTau40-k16 | 3.90 | 0.15 | 7.4 | SAXS | Mylonas *et al.*[11] |
| hTau40-k18 | 3.80 | 0.15 | 7.4 | SAXS | Mylonas *et al.*[11] |
| hTau40-k32 | 4.20 | 0.15 | 7.4 | SAXS | Mylonas *et al.*[11] |
| ibb | 2.52 | - | - | FRET | Fuertes *et al.*[12] |
| in | 2.16 | 0.05 | - | FRET | Hofmann *et al.*[9] |
| n49 | 1.37 | - | - | FRET | Fuertes *et al.*[12] |
| nls | 1.63 | - | - | FRET | Fuertes *et al.*[12] |
| nul | 2.66 | - | - | FRET | Fuertes *et al.*[12] |
| nus | 2.41 | - | - | FRET | Fuertes *et al.*[12] |
| p53 | 2.87 | - | - | SAXS | Wells *et al.*[13] |
| ProT$\alpha$-C | 3.00 | 0.04 | - | FRET | Muller-Spath *et al.*[5] |
| ProT$\alpha$-N | 2.55 | 0.04 | - | FRET | Muller-Spath *et al.*[5] |
| Protein G | 2.30 | - | - | SAXS | Smith *et al.*[14] |
| Protein L | 1.65 | 0.13 | - | FRET | Sherman & Haran [15] |
| R15 | 2.33 | 0.13 | - | FRET | Hofmann *et al.*[9] |
| R17 | 2.37 | 0.13 | - | FRET | Hofmann *et al.*[9] |
| RNaseA | 3.36 | 0.15 | 7.5 | SAXS | Riback *et al.*[8] |
| SH4 | 2.82 | 0.20 | 8.0 | SAXS | Arbesu *et al.*[16] |
| Sic1 | 3.00 | 0.20 | 7.5 | SAXS | Mittag *et al.*[17] |
| sNase | 2.12 | 0.02 | - | SAXS | Flanagan *et al.*[18] |
| ubiquitin | 2.52 | - | - | SAXS | Kohn *et al.*[2] |
| ul11 | 2.43 | - | - | SAXS | Metrick *et al.*[19] |
| yesg2 | 0.79 | 0.20 | - | SAXS | Ohnishi *et al.*[3] |
| yesg6 | 0.91 | 0.20 | - | SAXS | Ohnishi *et al.*[3] |

Solution condition and experimental method listed where available.

#### Table S4. Sequences of disordered proteins in the test set.

| | |
|---|---|
| Coint | MGSNGADNAH NNAFGGGKNP GIGNTSGAGS NGSASSNRGN SNGWSWSNKP HKNDGFHSDG SYHITFHGDN NSKPKPGGNS GNRGNNGDGA SSHHHHHH |
| (Hist5)2 | DSHAKRHHGY KRKFHEKHHS HRGYDSHAKR HHGYKRKFHE KHHSHRGY |
| hNL3cyt | MYRKDKRRQE PLRQPSPQRG AWAPELGAAP EEELAALQLG PTHHECEAGP PHDTLRLTAL PDYTLTLRRS PDDIPLMTPN TITMIPNSLV GLQTLHPYNT FAAGFNSTGL PHSHSTTRV |
| p15paf | MVRTKADSVP GTYRKVVAAR APRKVLGSST SATNSTSVSS RKAENKYAGG NPVCVRPTPK WQKGIGEFFR LSPKDSEKEN QIPEEAGSSG LGKAKRKACP LQPDHTNDEK E |
| A1 | GSMASASSSQ RGRSGSGNFG GGRGGGFGGN DNFGRGGNFS GRGGFGGSRG GGGYGGSGDG YNGFGNDGSN FGGGGSYNDF GNYNNQSSNF GPMKGGNFGG RSSGGSGGGG QYFAKPRNQG GYGGSSSSSS YGSGRRF |
| A1 -10F+7R+12D | GSMASADSSQ RDRDDRGNFG DGRGGGGGGN DNFGRGGNGS DRGGGGGSRG DGRYGGDGDR YNGGGNDGRN GGGGGSYNDG GNYNNQSSNG DPMKGGNGRD RSSGPYDRGG QYGAKPRNQG GYGGSSSSRS YGSDRRG |
| A1 -10R | GSMASASSSQ GGSSGSGNFG GGGGGGFGGN DNFGGGGNFS GSGGFGGSGG GGGYGGSGDG YNGFGNDGSN FGGGGSYNDF GNYNNQSSNF GPMKGGNFGG SSSGPYGGGG QYFAKPGNQG GYGGSSSSSS YGSGGGF |
| A1 -10R+10K | GSMASASSSQ KGKSGSGNFG GGKGGGFGGN DNFGKGGNFS GKGGFGGSKG GGGYGGSGDG YNGFGNDGSN FGGGGSYNDF GNYNNQSSNF GPMKGGNFGG KSSGGSGGGG QYFAKPKNQG GYGGSSSSSS YGSGKKF |
| A1 -12F+12Y | GSMASASSSQ RGRSGSGNYG GGRGGGYGGN DNYGRGGNYS GRGGYGGSRG GGGYGGSGDG YNGYGNDGSN YGGGGSYNDY GNYNNQSSNY GPMKGGNYGG RSSGGSGGGG QYYAKPRNQG GYGGSSSSSS YGSGRRY |
| A1 -12F+12Y-10R | GSMASASSSQ GGSSGSGNYG GGGGGGYGGN DNYGGGGNYS GSGGYGGSGG GGGYGGSGDG YNGYGNDGSN YGGGGSYNDY GNYNNQSSNY GPMKGGNYGG SSSGPYGGGG QYYAKPGNQG GYGGSSSSSS YGSGGGY |
| A1 -3R+3K | GSMASASSSQ RGKSGSGNFG GGRGGGFGGN DNFGRGGNFS GRGGFGGSKG GGGYGGSGDG YNGFGNDGSN FGGGGSYNDF GNYNNQSSNF GPMKGGNFGG RSSGGSGGGG QYFAKPRNQG GYGGSSSSSS YGSGRKF |
| A1 -4D | GSMASASSSQ RGRSGSGNFG GGRGGGFGGN GNFGRGGNFS GRGGFGGSRG GGGYGGSGGG YNGFGNSGSN FGGGGSYNGF GNYNNQSSNF GPMKGGNFGG RSSGPYGGGG QYFAKPRNQG GYGGSSSSSS YGSGRRF |
| A1 -6R | GSMASASSSQ GGRSGSGNFG GGRGGGFGGN DNFGGGGNFS GSGGFGGSRG GGGYGGSGDG YNGFGNDGSN FGGGGSYNDF GNYNNQSSNF GPMKGGNFGG SSSGPYGGGG QYFAKPGNQG GYGGSSSSSS YGSGGRF |
| A1 -6R+6K | GSMASASSSQ KGKSGSGNFG GGRGGGFGGN DNFGKGGNFS GRGGFGGSKG GGGYGGSGDG YNGFGNDGSN FGGGGSYNDF GNYNNQSSNF GPMKGGNFGG KSSGGSGGGG QYFAKPRNQG GYGGSSSSSS YGSGRKF |
| A1 -8F+4Y | GSMASASSSQ RGRSGSGNFG GGRGGGYGGN DNGGRGGNYS GRGGFGGSRG GGGYGGSGDG YNGGGNDGSN YGGGGSYNDS GNYNNQSSNF GPMKGGNYGG RSSGGSGGGG QYGAKPRNQG GYGGSSSSSS YGSGRRF |
| A1 -9F+3Y | GSMASASSSQ RGRSGSGNFG GGRGGGYGGN DNGGRGGNYS GRGGFGGSRG GGGYGGSGDG YNGGGNDGSN YGGGGSYNDS GNGNNQSSNF GPMKGGNYGG RSSGGSGGGG QYGAKPRNQG GYGGSSSSSS YGSGRRS |
| A1 -9F+6Y | GSMASASSSQ RGRSGSGNFG GGRGGGYGGN DNYGRGGNYS GRGGFGGSRG GGGYGGSGDG YNGGGNDGSN YGGGGSYNDS GNYNNQSSNF GPMKGGNYGG RSSGGSGGGG QYGAKPRNQG GYGGSSSSSS YGSGRRY |
| A1 +12D | GSMASADSSQ RDRDDSGNFG DGRGGGFGGN DNFGRGGNFS DRGGFGGSRG DGGYGGDGDG YNGFGNDGSN FGGGGSYNDF GNYNNQSSNF DPMKGGNFGD RSSGPYDGGG QYFAKPRNQG GYGGSSSSSS YGSDRRF |

| | |
|---|---|
| A1 +12E | GSMASAESSQ REREESGNFG EGRGGGFGGN DNFGRGGNFS ERGGFGGSRG EGGYGGEGDG YNGFGNDGSN FGGGGSYNDF GNYNNQSSNF EPMKGGNFGE RSSGPYEGGG QYFAKPRNQG GYGGSSSSSS YGSERRF |
| A1 +2R | GSMASASSSQ RGRSGSGNFG GGRGGGFGGN DNFGRGGNFS GRGGFGGSRG GGGYGGSGDG YNGFRNDGSN FGGGGRYNDF GNYNNQSSNF GPMKGGNFGG RSSGPYGGGG QYFAKPRNQG GYGGSSSSSS YGSGRRF |
| A1 +4D | GSMASASSSQ RDRSGSGNFG GGRGGGFGGN DNFGRGGNFS GRGDFGGSRG GGGYGGSGDG YNGFGNDGSN FGGGGSYNDF GNYNNQSSNF GPMKGGNFGG RSSDPYGGGG QYFAKPRNQG GYGGSSSSSS YDSGRRF |
| A1 +7F-7Y | GSMASASSSQ RGRSGSGNFG GGRGGGFGGN DNFGRGGNFS GRGGFGGSRG GGGFGGSGDG FNGFGNDGSN FGGGGSFNDF GNFNNQSSNF GPMKGGNFGG RSSGGSGGGG QFFAKPRNQG GFGGSSSSSS FGSGRRF |
| A1 +7K+12D | GSMASADSSQ RDRDDKGNFG DGRGGGFGGN DNFGRGGNFS DRGGFGGSRG DGKYGGDGDK YNGFGNDGKN FGGGGSYNDF GNYNNQSSNF DPMKGGNFKD RSSGPYDKGG QYFAKPRNQG GYGGSSSSKS YGSDRRF |
| A1 +7K+12D (b) | GSMASAKSSQ RDRDDDGNFG KGRGGGFGGN KNFGRGGNFS KRGGFGGSRG KGKYGGKGDD YNGFGNDGDN FGGGGSYNDF GNYNNQSSNF DPMDGGNFDD RSSGPYDDGG QYFADPRNQG GYGGSSSSKS YGSKRRF |
| A1 +7R | GSMASASSSQ RGRSGRGNFG GGRGGGFGGN DNFGRGGNFS GRGGFGGSRG GGRYGGSGDR YNGFGNDGRN FGGGGSYNDF GNYNNQSSNF GPMKGGNFRG RSSGPYGRGG QYFAKPRNQG GYGGSSSSRS YGSGRRF |
| A1 +8D | GSMASASSSQ RDRSGSGNFG GGRDGGFGGN DNFGRGDNFS GRGDFGGSRD GGGYGGSGDG YNGFGNDGSN FGGGGSYNDF GNYNNQSSNF GPMKGGNFGG RSSDPYGGGG QYFAKPRNQD GYGGSSSSSS YDSGRRF |

#### Table S5. Radii of gyration of proteins in the test set.

| protein | Rg (nm) | [ion] (M) | pH | Method | Reference |
|---|---|---|---|---|---|
| Coint | 2.83 | 0.40 | 7.6 | SAXS | Johnson *et al.*[20] |
| (Hist5)2 | 1.87 | 0.15 | 7.0 | SAXS | Fagerberg *et al.*[21] |
| hNL3cyt | 3.15 | 0.15 | 8.5 | SAXS | Paz *et al.*[22] |
| p15paf | 2.81 | 0.15 | 7.0 | SAXS | de Biasio *et al.*[23] |
| A1 | 2.76 | 0.15 | 7.0 | SAXS | Bremer *et al.*[24] |
| A1 -10F+7R+12D | 2.86 | 0.15 | 7.0 | SAXS | Bremer *et al.*[24] |
| A1 -10R | 2.67 | 0.15 | 7.0 | SAXS | Bremer *et al.*[24] |
| A1 -10R+10K | 2.85 | 0.15 | 7.0 | SAXS | Bremer *et al.*[24] |
| A1 -12F+12Y | 2.60 | 0.15 | 7.0 | SAXS | Bremer *et al.*[24] |
| A1 -12F+12Y-10R | 2.61 | 0.15 | 7.0 | SAXS | Bremer *et al.*[24] |
| A1 -3R+3K | 2.63 | 0.15 | 7.0 | SAXS | Bremer *et al.*[24] |
| A1 -4D | 2.64 | 0.15 | 7.0 | SAXS | Bremer *et al.*[24] |
| A1 -6R | 2.57 | 0.15 | 7.0 | SAXS | Bremer *et al.*[24] |
| A1 -6R+6K | 2.79 | 0.15 | 7.0 | SAXS | Bremer *et al.*[24] |
| A1 -8F+4Y | 2.71 | 0.15 | 7.0 | SAXS | Bremer *et al.*[24] |
| A1 -9F+3Y | 2.68 | 0.15 | 7.0 | SAXS | Bremer *et al.*[24] |
| A1 -9F+6Y | 2.66 | 0.15 | 7.0 | SAXS | Bremer *et al.*[24] |
| A1 +12D | 2.80 | 0.15 | 7.0 | SAXS | Bremer *et al.*[24] |
| A1 +12E | 2.85 | 0.15 | 7.0 | SAXS | Bremer *et al.*[24] |
| A1 +2R | 2.62 | 0.15 | 7.0 | SAXS | Bremer *et al.*[24] |
| A1 +4D | 2.72 | 0.15 | 7.0 | SAXS | Bremer *et al.*[24] |
| A1 +7F-7Y | 2.72 | 0.15 | 7.0 | SAXS | Bremer *et al.*[24] |
| A1 +7K+12D | 2.92 | 0.15 | 7.0 | SAXS | Bremer *et al.*[24] |
| A1 +7K+12D (b) | 2.56 | 0.15 | 7.0 | SAXS | Bremer *et al.*[24] |
| A1 +7R | 2.71 | 0.15 | 7.0 | SAXS | Bremer *et al.*[24] |
| A1 +8D | 2.69 | 0.15 | 7.0 | SAXS | Bremer *et al.*[24] |

Solution condition and experimental method listed where available.

#### Table S6. RNA experimental properties.

| Property | polyAde-30 | polyUra-30 | polyUra-40 | Reference |
|---|---|---|---|---|
| $R_g$, 20 mM NaCl (nm) | 2.72 | 3.06 | - | Plumridge *et al.*[25] |
| $R_g$, 100 mM NaCl (nm) | 2.45 | 2.68 | - | Plumridge *et al.*[25] |
| $R_g$, 200 mM NaCl (nm) | 2.51 | 2.51 | - | Plumridge *et al.*[25] |
| $R_g$, 400 mM NaCl (nm) | 2.38 | 2.33 | - | Plumridge *et al.*[25] |
| $R_g$, 600 mM NaCl (nm) | 2.22 | 2.33 | - | Plumridge *et al.*[25] |
| end-to-end, 25 mM NaCl (nm) | - | - | 6.86 | Chen *et al.*[26] |
| end-to-end, 50 mM NaCl (nm) | - | - | 6.75 | Chen *et al.*[26] |
| end-to-end, 100 mM NaCl (nm) | - | - | 6.63 | Chen *et al.*[26] |
| end-to-end, 200 mM NaCl (nm) | - | - | 6.42 | Chen *et al.*[26] |
| end-to-end, 400 mM NaCl (nm) | - | - | 6.26 | Chen *et al.*[26] |
| end-to-end, 800 mM NaCl (nm) | - | - | 6.04 | Chen *et al.*[26] |
| $P_{length}$, 20 mM NaCl (nm) | - | - | 2.25 | Chen *et al.*[26] |
| $P_{length}$, 40 mM NaCl (nm) | - | - | 2.13 | Chen *et al.*[26] |
| $P_{length}$, 80 mM NaCl (nm) | - | - | 1.98 | Chen *et al.*[26] |
| $P_{length}$, 150 mM NaCl (nm) | - | - | 1.76 | Chen *et al.*[26] |
| $P_{length}$, 275 mM NaCl (nm) | - | - | 1.60 | Chen *et al.*[26] |
| $P_{length}$, 500 mM NaCl (nm) | - | - | 1.40 | Chen *et al.*[26] |
| OCF \|i-j\| = 1 | 0.72 | 0.53 | - | Plumridge *et al.*[25] |
| OCF \|i-j\| = 2 | 0.35 | 0.34 | - | Plumridge *et al.*[25] |
| OCF \|i-j\| = 3 | 0.05 | 0.23 | - | Plumridge *et al.*[25] |
| OCF \|i-j\| = 4 | -0.11 | 0.18 | - | Plumridge *et al.*[25] |
| OCF \|i-j\| = 5 | -0.11 | 0.15 | - | Plumridge *et al.*[25] |
| OCF \|i-j\| = 6 | -0.02 | 0.11 | - | Plumridge *et al.*[25] |
| OCF \|i-j\| = 7 | 0.13 | 0.10 | - | Plumridge *et al.*[25] |
| OCF \|i-j\| = 8 | 0.23 | 0.10 | - | Plumridge *et al.*[25] |
| OCF \|i-j\| = 9 | 0.27 | 0.11 | - | Plumridge *et al.*[25] |
| OCF \|i-j\| = 10 | 0.22 | 0.14 | - | Plumridge *et al.*[25] |

polyAde-30
AAAAAAAAAA AAAAAAAAAA AAAAAAAAAA
polyUra-30
UUUUUUUUUU UUUUUUUUUU UUUUUUUUUU
polyUra-40
UUUUUUUUUU UUUUUUUUUU UUUUUUUUUU UUUUUUUUUU

OCF, orientational correlation factor
$OCF(|i-j|) = \langle cos\theta_{i,j} \rangle = \langle \hat{r}_i \cdot \hat{r}_j \rangle$, where $\hat{r}_i$ and $\hat{r}_j$ are normalized vectors between any $i, i+1$ and $j, j+1$ bonded residues in the chain, respectively.

#### Table S7. Homotypic protein phase separation systems composition.

| system | N chains | conc (mM) | conc (mg/mL) | box (nm) | Temp (K) | LLPS | LLPS Reference |
|---|---|---|---|---|---|---|---|
| FUS LCD | 72 | 0.120 | 2.140 | 100 | 298 | No | Kaur *et al.*[27] |
| FUS LCD | 90 | 0.150 | 2.670 | 100 | 298 | No | Kaur *et al.*[27] |
| FUS LCD | 105 | 0.175 | 3.110 | 100 | 298 | No | Kaur *et al.*[27] |
| FUS LCD | 120 | 0.200 | 3.550 | 100 | 298 | No | Kaur *et al.*[27] |
| FUS LCD | 132 | 0.220 | 3.910 | 100 | 298 | No | Kaur *et al.*[27] |
| FUS LCD | 150 | 0.250 | 4.440 | 100 | 298 | Yes | Kaur *et al.*[27] |
| FUS LCD | 157 | 0.260 | 4.620 | 100 | 298[(a)] | Yes | Kaur *et al.*[27] |
| FUS LCD | 181 | 0.300 | 5.340 | 100 | 298 | Yes | Kaur *et al.*[27] |
| FUS LCD | 240 | 0.400 | 7.090 | 100 | 298[(*)] | Yes | Kaur *et al.*[27] |
| FUS LCD | 362 | 0.600 | 10.680 | 100 | 298[(*)] | Yes | Kaur *et al.*[27] |
| FUS LCD | 432 | 0.720 | 12.760 | 100 | 298[(*)] | Yes | Kaur *et al.*[27] |
| LAF-1[RGG] | 36 | 0.018 | 0.300 | 150 | 298 | No | Elbaum-Garfinkle *et al.*[28] |
| LAF-1[RGG] | 42 | 0.021 | 0.350 | 150 | 298 | No | Elbaum-Garfinkle *et al.*[28] |
| LAF-1[RGG] | 47 | 0.023 | 0.390 | 150 | 298 | No | Elbaum-Garfinkle *et al.*[28] |
| LAF-1[RGG] | 51 | 0.025 | 0.420 | 150 | 298 | Yes | Elbaum-Garfinkle *et al.*[28] |
| LAF-1[RGG] | 54 | 0.027 | 0.450 | 150 | 298 | Yes | Elbaum-Garfinkle *et al.*[28] |
| LAF-1[RGG] | 60 | 0.030 | 0.500 | 150 | 298 | Yes | Elbaum-Garfinkle *et al.*[28] |
| LAF-1[RGG] | 72 | 0.036 | 0.600 | 150 | 298 | Yes | Elbaum-Garfinkle *et al.*[28] |
| LAF-1[RGG] | 166 | 0.276 | 4.620 | 100 | 298[(b)] | Yes | Elbaum-Garfinkle *et al.*[28] |
| A1 LCD | 49 | 0.010 | 0.120 | 200 | 298 | No | Bremer *et al.*[24] |
| A1 LCD | 90 | 0.020 | 0.240 | 200 | 298 | Yes | Bremer *et al.*[24] |
| A1 LCD | 135 | 0.030 | 0.366 | 200 | 298 | Yes | Bremer *et al.*[24] |
| A1 LCD | 180 | 0.040 | 0.488 | 200 | 298 | Yes | Bremer *et al.*[24] |
| A1 LCD | 213 | 0.353 | 4.620 | 100 | 298[(c)] | Yes | Bremer *et al.*[24] |
| hTau40-k18 | 96 | 0.020 | 0.274 | 200 | 298 | No | Ambadipudi *et al.*[29] |
| hTau40-k18 | 288 | 0.060 | 0.817 | 200 | 298 | Yes | Ambadipudi *et al.*[29] |
| hTau40-k18 | 336 | 0.070 | 0.955 | 200 | 298 | Yes | Ambadipudi *et al.*[29] |
| hTau40-k18 | 384 | 0.080 | 1.095 | 200 | 298 | Yes | Ambadipudi *et al.*[29] |
| hTau40-k18 | 203 | 0.338 | 4.620 | 100 | 298[(d)] | Yes | Ambadipudi *et al.*[29] |
| Ddx4 | 120 | 0.200 | 5.000 | 100 | 298[(e)] | Yes | Brady *et al.*[30] |

FUS LCD

MASNDYTQQA TQSYGAYPTQ PGQGYSQQSS QPYGQQSYSG YSQSTDTSGY GQSSYSSYGQ SQNSYGTQST PQGYGSTGGY GSSQSSQSSY GQQSSYPGYG QQPAPSSTSG SYGSSSQSSS YGQPQSGSYS QQPSYGGQQQ SYGQQQSYNP PQGYGQQNQY NSSSGGGGGG GGG

LAF-1[RGG]

MESNQSNNGG SGNAALNRGG RYVPPHLRGG DGGAAAAASA GGDDRRGGAG GGGYRRGGGN SGGGGGGGYD RGYNDNRDDR DNRGGSGGYG RDRNYEDRGY NGGGGGGGNR GYNNNRGGGG GGYNRQDRGD GGSSNFSRGG YNNRDEGSDN RGSGRSYNND RRDNGGDG

A1 LCD

GSMASASSSQ RGRSGSGNFG GGRGGGFGGN DNFGRGGNFS GRGGFGGSRG GGGYGGSGDG YNGFGNDGSN FGGGGSYNDF GNYNNQSSNF GPMKGGNFGG RSSGGSGGGG QYFAKPRNQG GYGGSSSSSS YGSGRRF

hTau40-k18

MQTAPVPMPD LKNVKSKIGS TENLKHQPGG GKVQIINKKL DLSNVQSKCG SKDNIKHVPG GGSVQIVYKP VDLSKVTSKC GSLGNIHHKP GGGQVEVKSE KLDFKDRVQS KIGSLDNITH VPGGGNKKIE

Ddx4

MGDEDWEAEI NPHMSSYVPI FEKDRYSGEN GDNFNRTPAS SSEMDDGPSR RDHFMKSGFA SGRNFGNRDA GECNKRDNTS TMGGFGVGKS FGNRGFSNSR FEDGDSSGFW RESSNDCEDN PTRNRGFSKR GGYRDGNNSE ASGPYRRGGR GSFRGCRGGF GLGSPNNDLD PDECMQRTGG LFGSRRPVLS GTGNGDTSQS RSGSGSERGG YKGLNEEVIT GSGKNSWKSE AEGGES

[(a)] also simulated at 260, 270, 280, 290, and 310 K.
[(b)] also simulated at 260, 270, 280, 290, 310, 320, and 334 K.
[(c)] also simulated at 260, 270, 280, 290, 310, 320, and 334 K.
[(d)] also simulated at 260, 270, 280, 290, 310, and 320 K.
[(e)] also simulated at 260, 270, 280, 290, 310, 320, 330, 340, 350, and 360 K.
[(*)] only simulated using Regy *et al.*[31] and Dannenhoffer *et al.*[32] force fields.

#### Table S8. Heterotypic protein phase separation systems composition.

| system | N chains | conc (mM) | conc (mg/mL) | box (nm) | Temp (K) | LLPS | LLPS Reference |
|---|---|---|---|---|---|---|---|
| $(RGRGG)_5$ / FUS LCD | 132/132 | 0.220/0.220 | 0.533/3.550 | 100 | 298 | No | Kaur *et al.*[27] |
| $(RGRGG)_5$ / FUS LCD | 396/132 | 0.660/0.220 | 1.598/3.550 | 100 | 298 | Yes | Kaur *et al.*[27] |
| $(RGRGG)_5$ / FUS LCD | 105/105 | 0.175/0.175 | 0.425/3.110 | 100 | 298 | No | Kaur *et al.*[27] |
| $(RGRGG)_5$ / FUS LCD | 210/105 | 0.350/0.175 | 0.850/3.110 | 100 | 298 | No | Kaur *et al.*[27] |
| $(RGRGG)_5$ / FUS LCD | 315/105 | 0.575/0.175 | 1.275/3.110 | 100 | 298 | Yes | Kaur *et al.*[27] |
| $(RGRGG)_5$ / FUS LCD | 420/105 | 0.700/0.175 | 1.694/3.110 | 100 | 298 | Yes | Kaur *et al.*[27] |
| $(RGRGG)_5$ / FUS LCD | 525/105 | 0.875/0.175 | 2.117/3.110 | 100 | 298 | Yes | Kaur *et al.*[27] |
| $(RGRGG)_5$ / FUS LCD | 840/105 | 1.400/0.175 | 3.385/3.110 | 100 | 298 | Yes | Kaur *et al.*[27] |
| $(RGRGG)_5$ / FUS LCD | 90/90 | 0.150/0.150 | 0.363/2.670 | 100 | 298 | No | Kaur *et al.*[27] |
| $(RGRGG)_5$ / FUS LCD | 450/90 | 0.750/0.150 | 1.814/2.670 | 100 | 298 | No | Kaur *et al.*[27] |
| $(RGRGG)_5$ / FUS LCD | 900/90 | 1.500/0.150 | 3.630/2.670 | 100 | 298 | Yes | Kaur *et al.*[27] |
| $(RGRGG)_5$ / FUS LCD | 990/90 | 1.643/0.150 | 3.990/2.670 | 100 | 298 | Yes | Kaur *et al.*[27] |
| $(RGRGG)_5$ / FUS LCD | 1080/90 | 1.794/0.150 | 4.355/2.670 | 100 | 298 | Yes | Kaur *et al.*[27] |
| $(RGRGG)_5$ / FUS LCD | 1350/90 | 2.242/0.150 | 5.444/2.670 | 100 | 298 | Yes | Kaur *et al.*[27] |
| $(RGRGG)_5$ / FUS LCD | 144/72 | 0.240/0.120 | 0.580/2.140 | 100 | 298 | No | Kaur *et al.*[27] |
| $(RGRGG)_5$ / FUS LCD | 720/72 | 1.200/0.120 | 2.903/2.140 | 100 | 298 | Yes | Kaur *et al.*[27] |
| $(RGRGG)_5$ / FUS LCD | 1080/72 | 1.800/0.120 | 4.355/2.140 | 100 | 298 | Yes | Kaur *et al.*[27] |
| FUS $LCD^{RGG3}$ / FUS LCD | 120/120 | 0.200/0.200 | 0.690/3.550 | 100 | 298 | No | Kaur *et al.*[27] |
| FUS $LCD^{RGG3}$ / FUS LCD | 240/120 | 0.400/0.200 | 1.380/3.550 | 100 | 298 | Yes | Kaur *et al.*[27] |
| FUS $LCD^{RGG3}$ / FUS LCD | 360/120 | 0.600/0.200 | 2.070/3.550 | 100 | 298 | Yes | Kaur *et al.*[27] |
| FUS $LCD^{RGG3}$ / FUS LCD | 105/105 | 0.175/0.175 | 0.605/3.110 | 100 | 298 | No | Kaur *et al.*[27] |
| FUS $LCD^{RGG3}$ / FUS LCD | 210/105 | 0.350/0.175 | 1.210/3.110 | 100 | 298 | No | Kaur *et al.*[27] |
| FUS $LCD^{RGG3}$ / FUS LCD | 315/105 | 0.525/0.175 | 1.813/3.110 | 100 | 298 | Yes | Kaur *et al.*[27] |
| FUS $LCD^{RGG3}$ / FUS LCD | 630/105 | 1.050/0.175 | 3.620/3.110 | 100 | 298 | Yes | Kaur *et al.*[27] |
| FUS $LCD^{RGG3}$ / FUS LCD | 90/90 | 0.150/0.150 | 0.520/2.670 | 100 | 298 | No | Kaur *et al.*[27] |
| FUS $LCD^{RGG3}$ / FUS LCD | 180/90 | 0.300/0.150 | 1.034/2.670 | 100 | 298 | No | Kaur *et al.*[27] |
| FUS $LCD^{RGG3}$ / FUS LCD | 270/90 | 0.450/0.150 | 1.550/2.670 | 100 | 298 | No | Kaur *et al.*[27] |
| FUS $LCD^{RGG3}$ / FUS LCD | 360/90 | 0.600/0.150 | 2.070/2.670 | 100 | 298 | Yes | Kaur *et al.*[27] |
| FUS $LCD^{RGG3}$ / FUS LCD | 450/90 | 0.750/0.150 | 2.585/2.670 | 100 | 298 | Yes | Kaur *et al.*[27] |
| FUS $LCD^{RGG3}$ / FUS LCD | 540/90 | 0.900/0.150 | 3.105/2.670 | 100 | 298 | Yes | Kaur *et al.*[27] |
| FUS $LCD^{RGG3}$ / FUS LCD | 144/72 | 0.240/0.120 | 0.830/2.140 | 100 | 298 | No | Kaur *et al.*[27] |
| FUS $LCD^{RGG3}$ / FUS LCD | 288/72 | 0.480/0.120 | 1.655/2.140 | 100 | 298 | No | Kaur *et al.*[27] |
| FUS $LCD^{RGG3}$ / FUS LCD | 432/72 | 0.720/0.120 | 2.485/2.140 | 100 | 298 | Yes | Kaur *et al.*[27] |
| FUS $LCD^{RGG3}$ / FUS LCD | 504/72 | 0.840/0.120 | 2.896/2.140 | 100 | 298 | Yes | Kaur *et al.*[27] |
| FUS $LCD^{RGG3}$ / FUS LCD | 576/72 | 0.960/0.120 | 3.312/2.140 | 100 | 298 | Yes | Kaur *et al.*[27] |

$(RGRGG)_5$

RGRGGRGRGG RGRGGRGRGG RGRGG

FUS $LCD^{RGG3}$

RRGGRGGYDR GGYRGRGGDR GGFRGGRGGG DRGC

#### Table S9. Protein – RNA phase separation systems composition.

| System | N chains | conc (mM) | conc (mg/mL) | box (nm) | Temp (K) | LLPS Reference |
|---|---|---|---|---|---|---|
| polyAde-21 / $(RRLR)_6$-SSSGSS | 126/147 | 0.210/0.240 | 1.390/0.975 | 100 | 298 | Bai *et al.*[33] |
| polyUra-40 / FUS $LCD^{RGG3}$ | 197/696 | 0.330/1.160 | 4.000/4.000 | 100 | 298 | Kaur *et al.*[27] |
| polyUra-10 / polyArg-50 | 360/72 | 0.600/0.120 | 1.825/0.945 | 100 | 298 | Fisher & Elbaum-Garfinkle[34] |
| polyAde-500 / $(RGRGG)_5$ | 30/1786 | 0.990/0.900 | 0.006/0.371 | 200 | 298 | Alshareedah *et al.*[35] |

$(RRLR)_6$-SSSGSS

RRLRRRLRRR LRRRLRRRLR RRLRSSSGSS

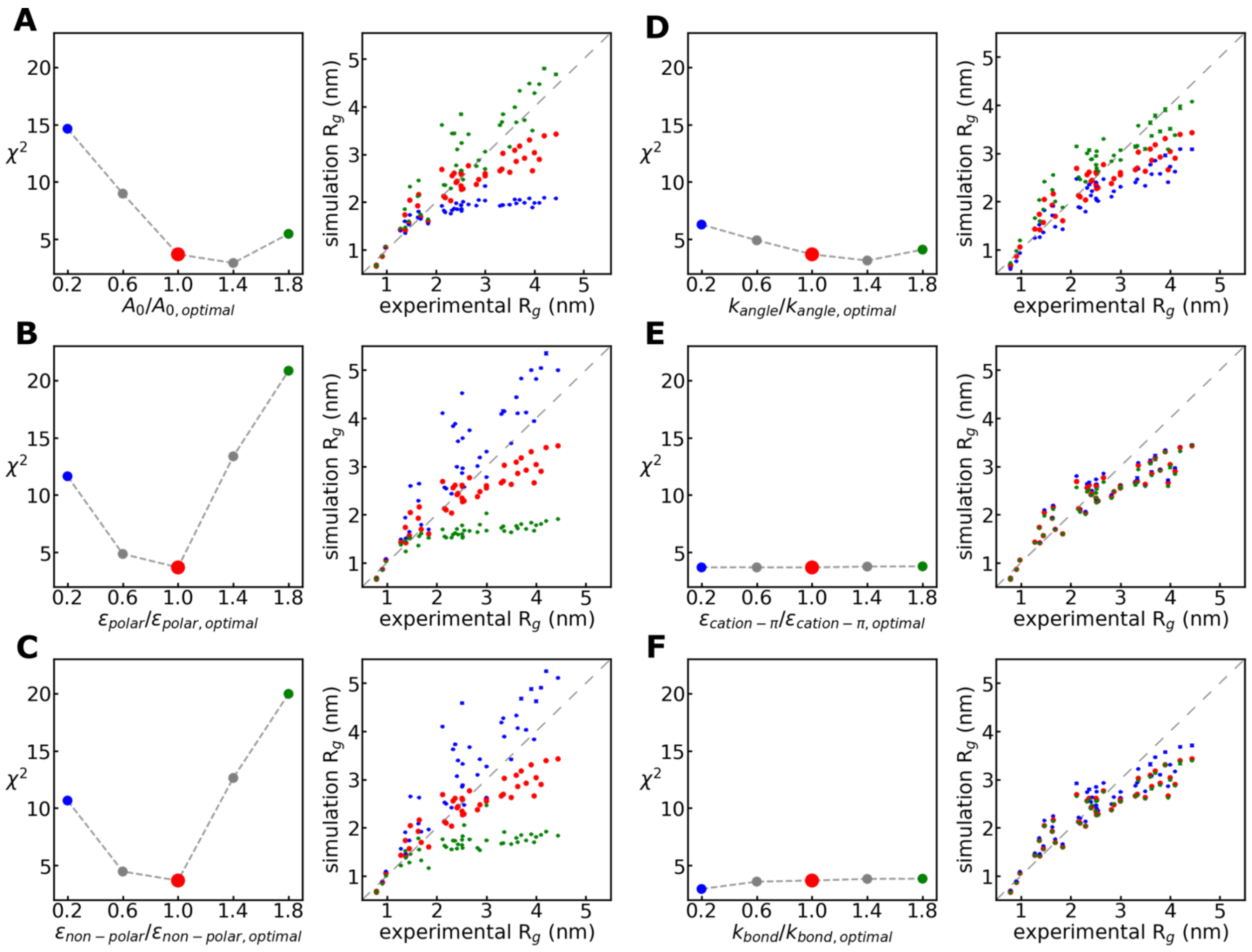


**Figure S1. Protein radius of gyration (Rg) sensitivity to model parameters.** The $\chi^2_{\exp\,vs\,MD}$ values are plotted as a function of the parameters value:value$_{\text{optimal}}$ ratio (left panels) for polar residues $A_0$ (A), $\varepsilon_{polar}$ (B), $\varepsilon_{non-polar}$ (C), $k_{angle}$ (D), $\varepsilon_{cation-\pi}$ (E), and $k_{bond}$ (F). Simulated vs experimental $R_g$ plots (right panels) are also shown for the lowest (blue), optimal (red), and highest (green) values evaluated for each parameter.

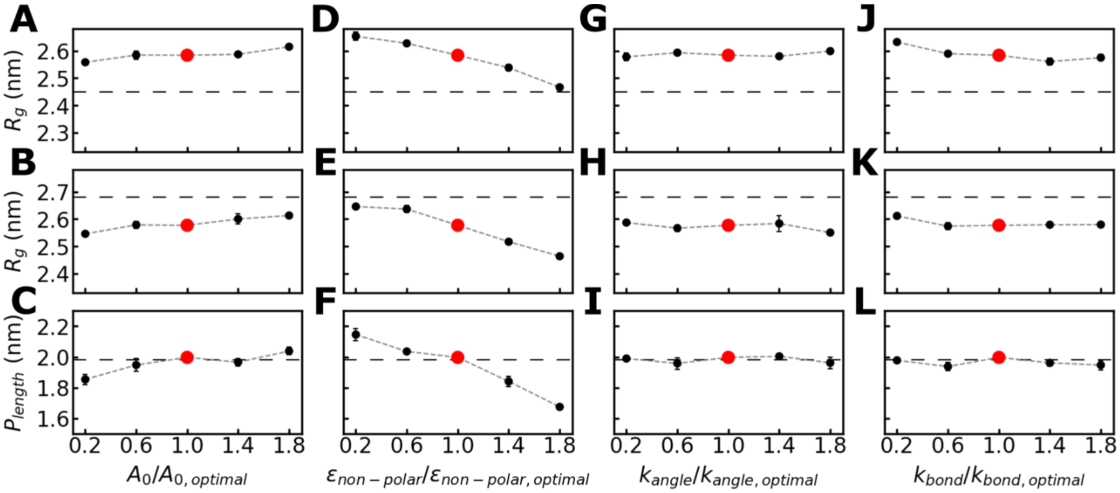


**Figure S2. RNA polymer properties sensitivity to model parameters.** Radius of gyration of polyAde-30 (A, D, and G) and polyUra-30 (B, E, and H), and persistence length ($P_{length}$) for polyUra-40 (C, F, and I) are plotted as a function of the parameters value:value$_{optimal}$ ratio for polar residues $A_0$ (A-C), $\varepsilon_{non-polar}$(D-E), $k_{angle}$ (G-I), and $k_{bond}$ (J-L) parameters. The data shown was obtained with $\kappa$ = 1 nm (equivalent to ~100 mM of salt). Experimental values for each case are shown as dashed lines [25, 26]. The optimal value used in COCOMO is highlighted in red.

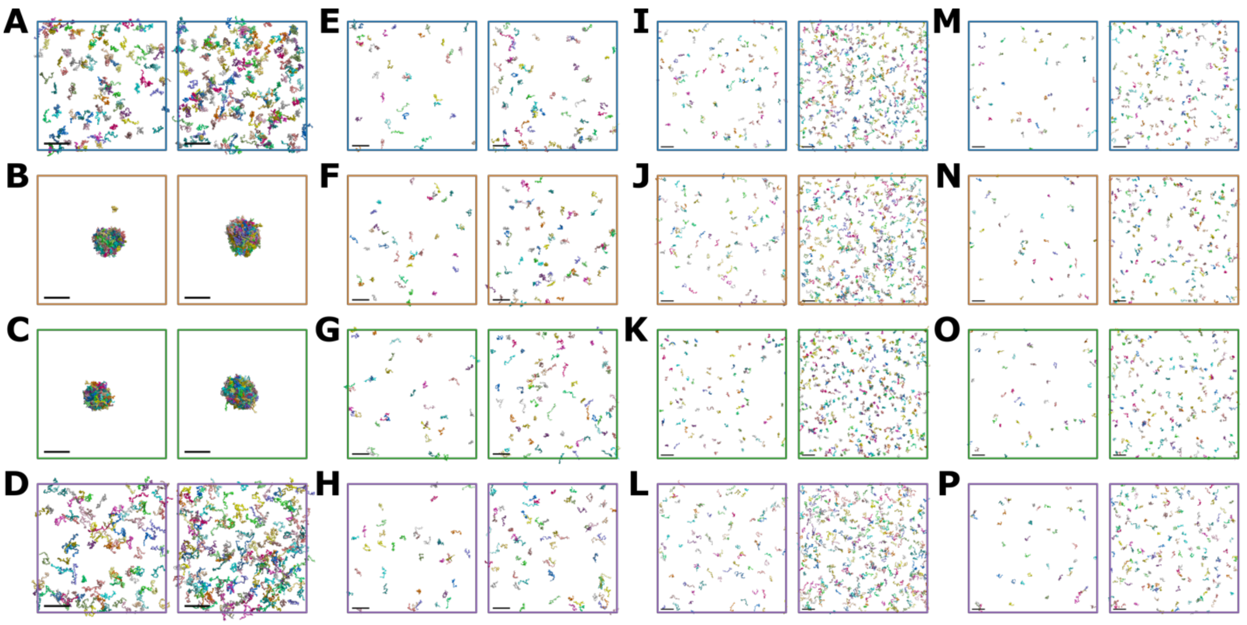

**Figure S3. Representative snapshots for homotypic protein systems simulated with different models.** Final frames are shown for systems at concentrations below (left) and above (right) the experimental thresholds for FUS LCD (A-D), LAF-1$^{RGG}$ (E-H), hTau40-k18 (I-L), and A1 LCD (M-P). Different rows correspond to different models: Regy *et al.* 2021[31] (A,E, I, and M; blue PBC cell), Tesei *et al.* 2021[36] (B, F, J, and N; tan PBC cell), Dignon *et al.* 2018[37] (C, G, K, and O; green PBC cell), and Dannenhoffer *et al.* 2021[32] (D, H, L, and P; purple PBC cell) [31, 32, 36, 37]. Colors are used to distinguish different chains. The size bar represents 20 nm. Table S4 gives the exact concentration values of each system.

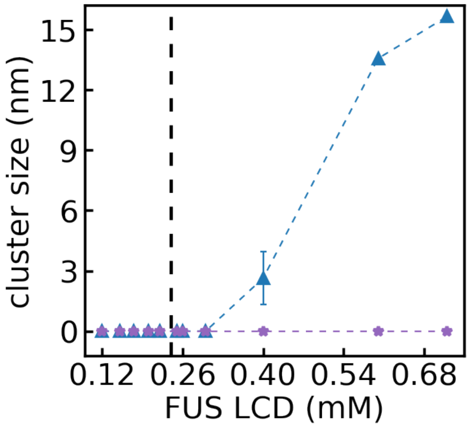


**Figure S4. Phase separation in FUS LCD with other models**. Cluster sizes as a function of concentration for FUS LCD using models from Regy *et al.* 2021[31] (blue triangle) and Dannenhoffer *et al.* 2021[38] (purple star) at higher concentrations. The experimental LLPS concentration threshold is shown as in **Figure 3**.

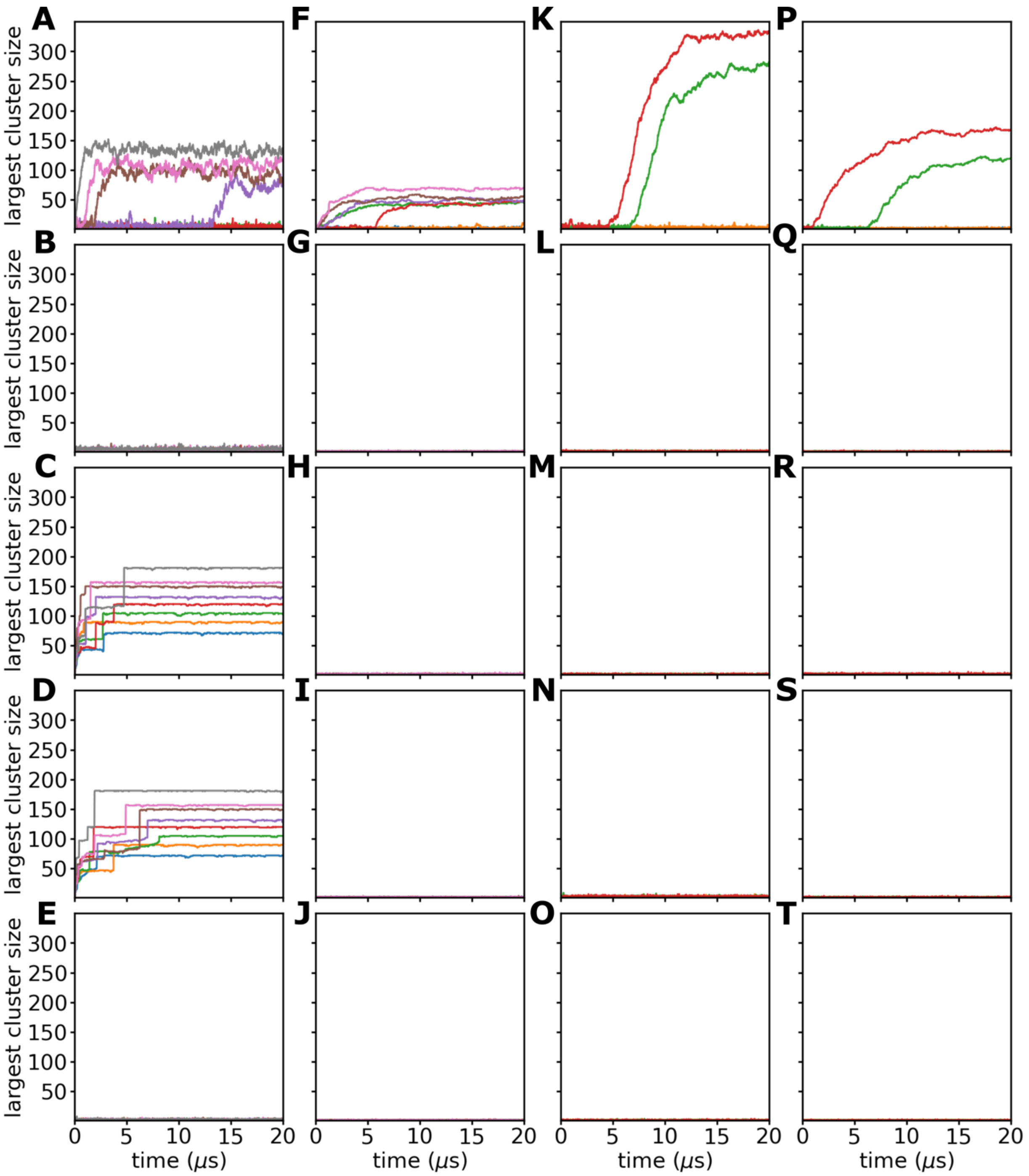


**Figure S5. Cluster formation during homotypic phase separation.** The size of the largest cluster is shown as a function of simulation time for FUS LCD (A-E), LAF-1$^{RGG}$ (F-J), hTau40-k18 (K-O), and A1 LCD (P-T). Results are shown row-wise for COCOMO (A, F, K, and P), the Regy *et al.* 2021[31] model (B, G, L, and Q), the Tesei *et al.* 2021[36] model (C, H, M, and R), the Dignon *et al.* 2018[37] model (D, I, N, and S), and the Dannenhoffer *et al.* 2021[32] model (E, J, O, and T). Each trace corresponds to a different concentration of the protein, colored in increasing order as blue, orange, green, red, purple, brown, pink, and gray (see **Table S4** for the concentration values of each system).

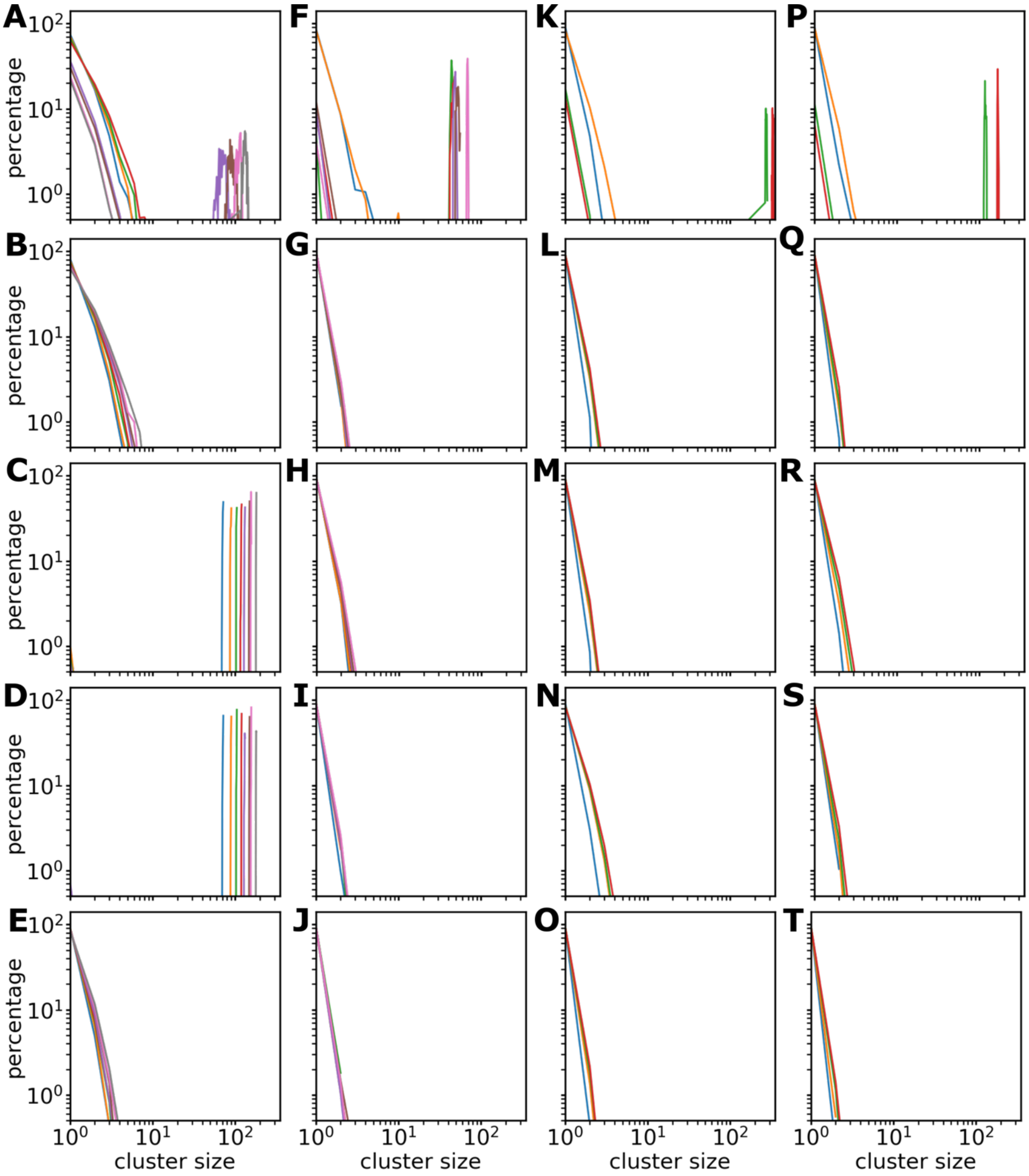


**Figure S6. Cluster size distributions in homotypic protein systems.** Cluster size distributions were obtained from simulation snapshots during the last 4 μs of the trajectories for FUS LCD (A-E), LAF-1$^{RGG}$ (F-J), hTau40-k18 (K-O), and A1 LCD (P-T). Results are shown row-wise for different models and colored as in **Figure S5**.

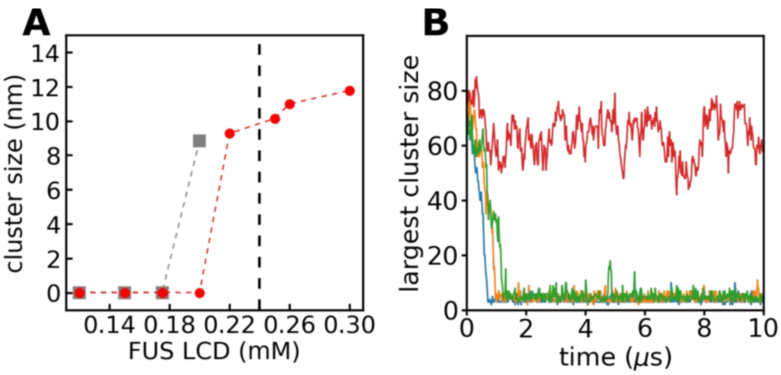


**Figure S7. Reversible phase separation in FUS LCD with COCOMO.** (A) Cluster sizes as a function of concentration for FUS LCD using COCOMO in simulations started from random orientations (red circles) and simulations starting from the condensate at 0.220 mM but with increased box sizes to achieve lower concentrations (grey squares). (B) The size of the largest cluster as a function of simulation time for FUS LCD for the simulations starting from the condensate at 0.220 mM with final concentration of 0.120 (blue), 0.150 (orange), 0.175 (green), and 0.200 mM (red).

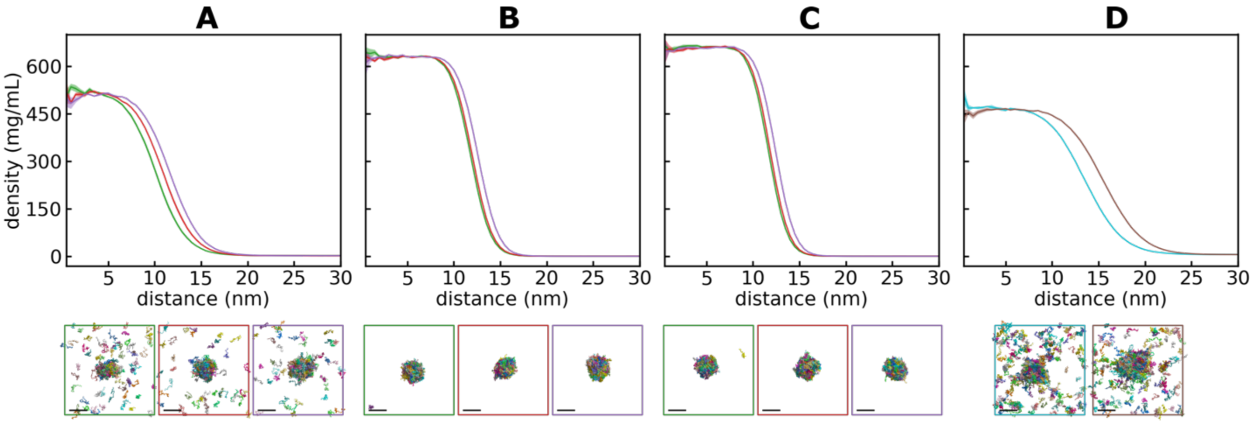


**Figure S8. Density profiles of FUS LCD simulation at concentration above the experimental LLPS threshold.** Results are shown for COCOMO (A), Tesei *et al.* 2021[36] (B), Dignon *et al.* 2018[37] (C), and Regy *et al.* 2021[31] (D) models. Each trace is the averaged profile for the last 4 μs of the simulation with standard error showed shaded. Initial FUS LCD concentrations were 0.25 (green), 0.26 (red), and 0.30 mM (purple) for A-C, as well as 0.6 (cyan) and 0.72 mM (brown) for D. The density was calculated as a function of distance starting from the center of mass of the droplet. Lower panels show snapshots at the end of each trajectory (PBC cell colors match concentration colors), with a size bar that represents 20 nm.

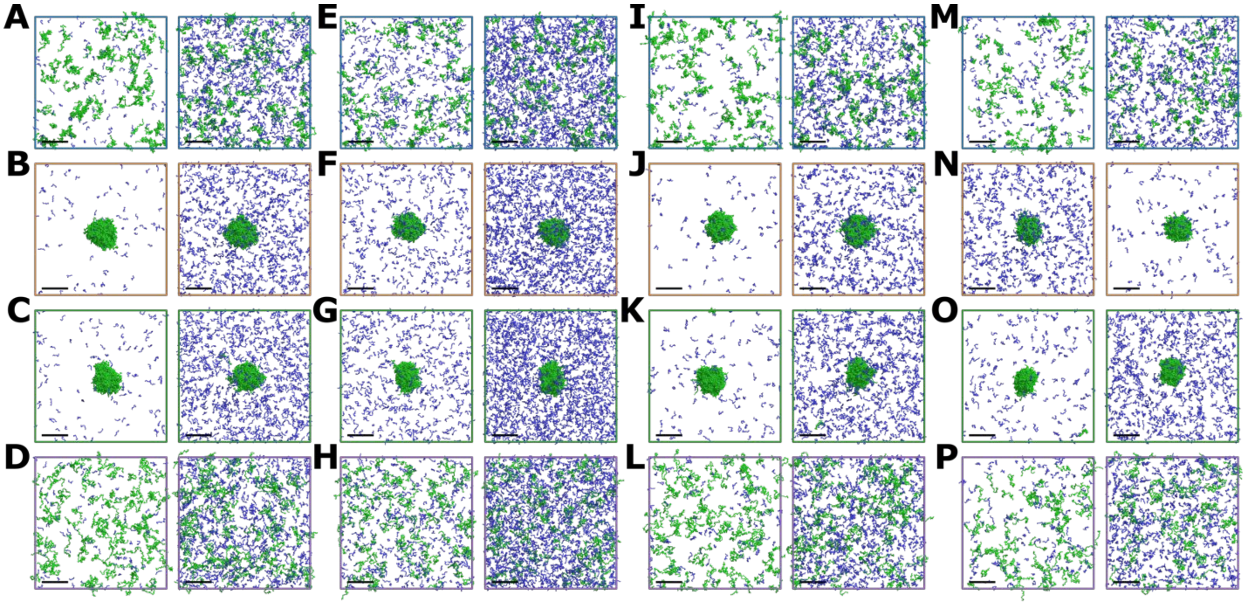


**Figure S9. Representative snapshots for heterotypic LLPS in different systems.** Final frames are shown for the lowest (left) and highest (right) concentrations tried for the protein system of FUS LCD 0.175 mM with $[RGRGG]_5$ peptide (A-D), FUS LCD 0.150 mM with $[RGRGG]_5$ peptide (E-H), FUS LCD 0.175 mM with FUS LCD$^{RGG3}$ peptide (I-L), and FUS LCD 0.120 mM with FUS LCD$^{RGG3}$ peptide (M-P). Results are shown row-wise for different models: Regy *et al.* 2021[31] (A,E, I, and M; blue PBC cell), Tesei *et al.* 2021[36] (B, F, J, and N; tan PBC cell), Dignon *et al.* 2018[37] (C, G, K, and O; green PBC cell), and Dannenhoffer *et al.* 2021[32] (D, H, L, and P; purple PBC cell) [31, 32, 36, 37]. The FUS LCD and peptide chains are colored in green and blue, respectively. The size bar represents 20 nm. Refer to table S4 for the exact concentration values of each system.

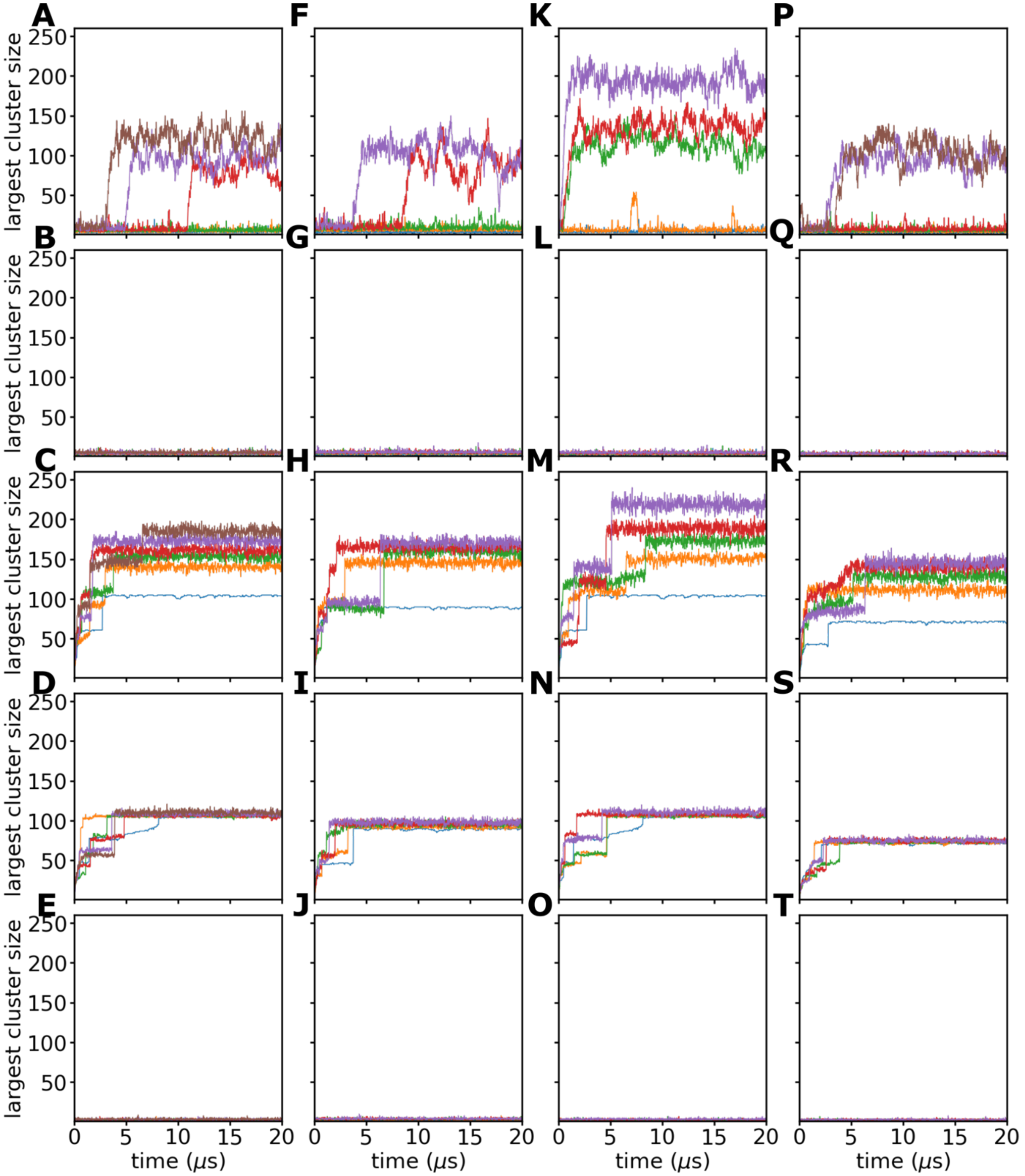


**Figure S10. Cluster formation in heterotypic phase separation.** The time evolution of the largest cluster size is shown as in **Figure S4** for FUS LCD 0.175 mM with [RGRGG]$_5$ peptide (A-E), FUS LCD 0.150 mM with [RGRGG]$_5$ peptide (F-J), FUS LCD 0.175 mM with FUS LCD$^{RGG3}$ peptide (K-O), and FUS LCD 0.120 mM with FUS LCD$^{RGG3}$ peptide (P-T). Each trace corresponds to a different concentration of peptide added, colored in increasing order as blue, orange, green, red, purple, and brown. Results are shown row-wise for different models as in Fig S4.

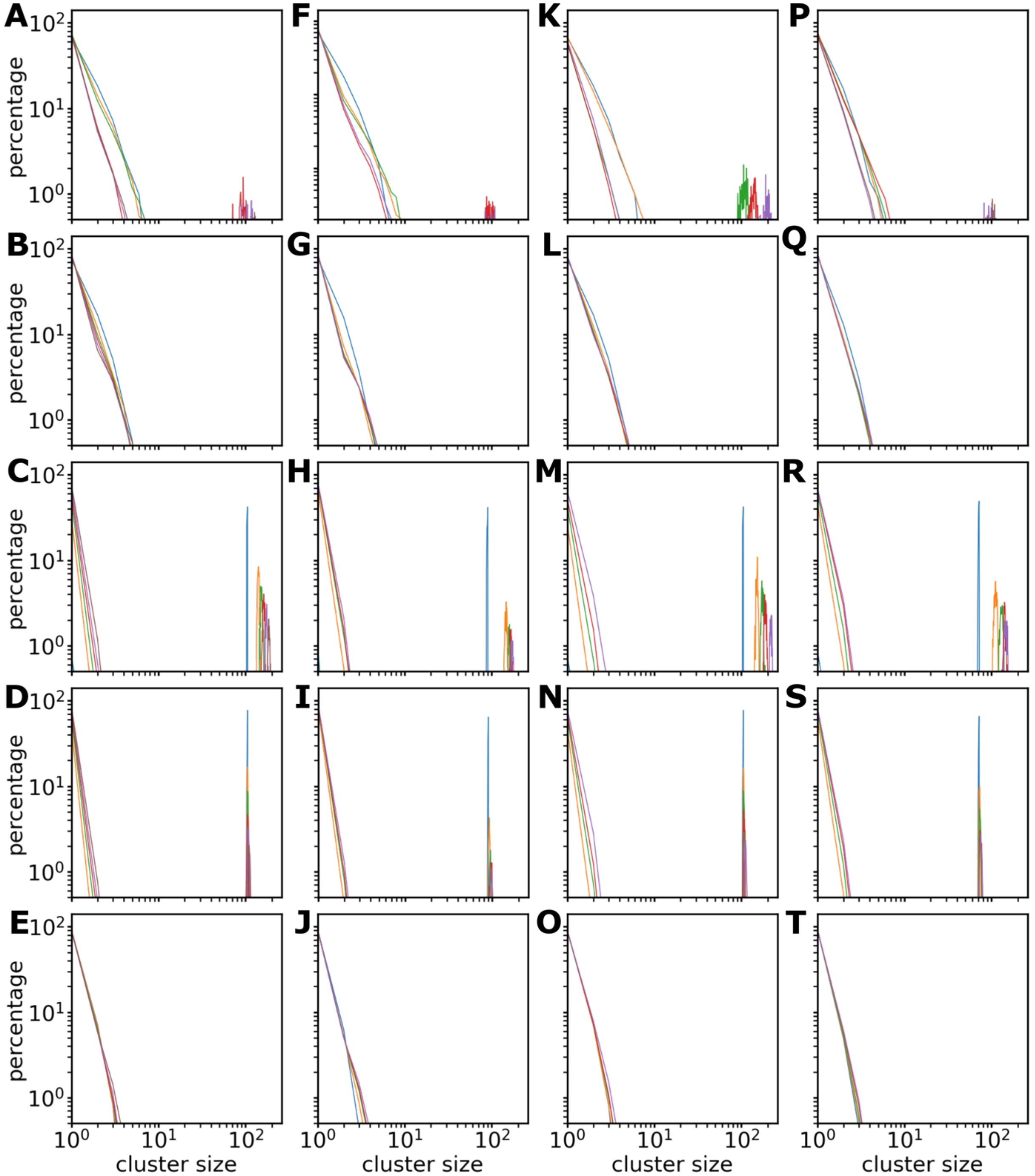


**Figure S11. Cluster size distributions in heterotypic phase separation.** Results are shown as in **Figure S5** for FUS LCD 0.175 mM with $[RGRGG]_5$ peptide (A-E), FUS LCD 0.150 mM with $[RGRGG]_5$ peptide (F-J), FUS LCD 0.175 mM with FUS $LCD^{RGG3}$ peptide (K-O), and FUS LCD 0.120 mM with FUS $LCD^{RGG3}$ peptide (P-T). Results are shown row-wise for different models and colored as in **Figure S10**.

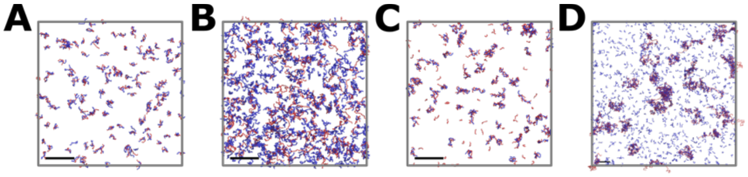


**Figure S12. Representative conformations of different RNA-protein systems.** Final frames of simulations using the Regy 2020 model[39] in the mixes of polyAde-21 – $(RRLR)_6$-SSSGSS (A), polyUra-40 – FUS LCD$^{RGG3}$ (B), polyUra-10 – polyArg-50 (C), and polyAde-500 – $(RGRGG)_5$ (D). RNA and protein chains were colored in red and blue, respectively. The size bar represents 20 nm.

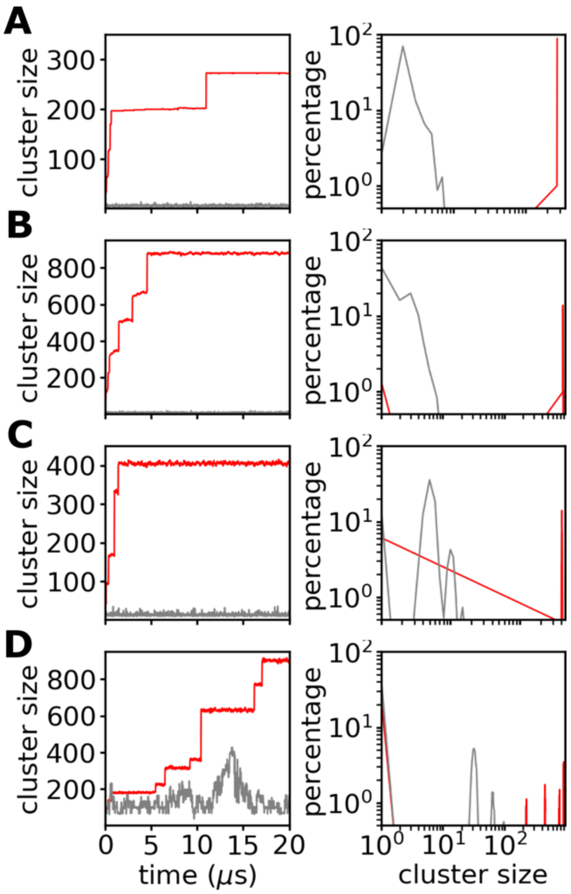


**Figure S13. Cluster characterization in RNA – protein phase separation.** Size of the largest cluster *vs* simulation time and cluster size distributions during the last 4 μs of the trajectory are showed for our model (red) and Regy 2020 (gray)[39] using polyAde-21 – $(RRLR)_6$-SSSGSS (A), polyUra-40 – FUS $LCD^{RGG3}$ (B), polyUra-10 – polyArg-50 (C), and polyAde-500 – $(RGRGG)_5$ (D).

#### SUPPLEMENTARY REFERENCES


(1) Kjaergaard, M.; Norholm, A. B.; Hendus-Altenburger, R.; Pedersen, S. F.; Poulsen, F. M.; Kragelund, B. B. Temperature-dependent structural changes in intrinsically disordered proteins: formation of alpha-helices or loss of polyproline II? *Protein Sci* **2010**, *19*, 1555-1564.
(2) Kohn, J. E.; Millett, I. S.; Jacob, J.; Zagrovic, B.; Dillon, T. M.; Cingel, N.; Dothager, R. S.; Seifert, S.; Thiyagarajan, P.; Sosnick, T. R.; et al. Random-coil behavior and the dimensions of chemically unfolded proteins. *Proc Natl Acad Sci U S A* **2004**, *101*, 12491-12496.
(3) Ohnishi, S.; Kamikubo, H.; Onitsuka, M.; Kataoka, M.; Shortle, D. Conformational preference of polyglycine in solution to elongated structure. *J Am Chem Soc* **2006**, *128*, 16338-16344.
(4) Nath, A.; Sammalkorpi, M.; DeWitt, D. C.; Trexler, A. J.; Elbaum-Garfinkle, S.; O'Hern, C. S.; Rhoades, E. The conformational ensembles of alpha-synuclein and tau: combining single-molecule FRET and simulations. *Biophys J* **2012**, *103*, 1940-1949.
(5) Muller-Spath, S.; Soranno, A.; Hirschfeld, V.; Hofmann, H.; Ruegger, S.; Reymond, L.; Nettels, D.; Schuler, B. From the Cover: Charge interactions can dominate the dimensions of intrinsically disordered proteins. *Proc Natl Acad Sci U S A* **2010**, *107*, 14609-14614.
(6) Choy, W. Y.; Mulder, F. A.; Crowhurst, K. A.; Muhandiram, D. R.; Millett, I. S.; Doniach, S.; Forman-Kay, J. D.; Kay, L. E. Distribution of molecular size within an unfolded state ensemble using small-angle X-ray scattering and pulse field gradient NMR techniques. *J Mol Biol* **2002**, *316*, 101-112.
(7) Lens, Z.; Dewitte, F.; Monte, D.; Baert, J. L.; Bompard, C.; Senechal, M.; Van Lint, C.; de Launoit, Y.; Villeret, V.; Verger, A. Solution structure of the N-terminal transactivation domain of ERM modified by SUMO-1. *Biochem Biophys Res Commun* **2010**, *399*, 104-110.
(8) Riback, J. A.; Bowman, M. A.; Zmyslowski, A. M.; Knoverek, C. R.; Jumper, J. M.; Hinshaw, J. R.; Kaye, E. B.; Freed, K. F.; Clark, P. L.; Sosnick, T. R. Innovative scattering analysis shows that hydrophobic disordered proteins are expanded in water. *Science* **2017**, *358*, 238-241.
(9) Hofmann, H.; Soranno, A.; Borgia, A.; Gast, K.; Nettels, D.; Schuler, B. Polymer scaling laws of unfolded and intrinsically disordered proteins quantified with single-molecule spectroscopy. *Proc Natl Acad Sci U S A* **2012**, *109*, 16155-16160.
(10) Cragnell, C.; Durand, D.; Cabane, B.; Skepo, M. Coarse-grained modeling of the intrinsically disordered protein Histatin 5 in solution: Monte Carlo simulations in combination with SAXS. *Proteins* **2016**, *84*, 777-791.
(11) Mylonas, E.; Hascher, A.; Bernado, P.; Blackledge, M.; Mandelkow, E.; Svergun, D. I. Domain conformation of tau protein studied by solution small-angle X-ray scattering. *Biochemistry* **2008**, *47*, 10345-10353.
(12) Fuertes, G.; Banterle, N.; Ruff, K. M.; Chowdhury, A.; Mercadante, D.; Koehler, C.; Kachala, M.; Estrada Girona, G.; Milles, S.; Mishra, A.; et al. Decoupling of size and shape fluctuations in heteropolymeric sequences reconciles discrepancies in SAXS vs. FRET measurements. *Proc Natl Acad Sci U S A* **2017**, *114*, E6342-E6351.
(13) Wells, M.; Tidow, H.; Rutherford, T. J.; Markwick, P.; Jensen, M. R.; Mylonas, E.; Svergun, D. I.; Blackledge, M.; Fersht, A. R. Structure of tumor suppressor p53 and its intrinsically disordered N-terminal transactivation domain. *Proc Natl Acad Sci U S A* **2008**, *105*, 5762-5767.
(14) Smith, C. K.; Bu, Z.; Anderson, K. S.; Sturtevant, J. M.; Engelman, D. M.; Regan, L. Surface point mutations that significantly alter the structure and stability of a protein's denatured state. *Protein Sci* **1996**, *5*, 2009-2019.
(15) Sherman, E.; Haran, G. Coil-globule transition in the denatured state of a small protein. *Proc Natl Acad Sci U S A* **2006**, *103*, 11539-11543.
(16) Arbesu, M.; Maffei, M.; Cordeiro, T. N.; Teixeira, J. M.; Perez, Y.; Bernado, P.; Roche, S.; Pons, M. The Unique Domain Forms a Fuzzy Intramolecular Complex in Src Family Kinases. *Structure* **2017**, *25*, 630-640 e634.
(17) Mittag, T.; Marsh, J.; Grishaev, A.; Orlicky, S.; Lin, H.; Sicheri, F.; Tyers, M.; Forman-Kay, J. D. Structure/function implications in a dynamic complex of the intrinsically disordered Sic1 with the Cdc4 subunit of an SCF ubiquitin ligase. *Structure* **2010**, *18*, 494-506.
(18) Flanagan, J. M.; Kataoka, M.; Shortle, D.; Engelman, D. M. Truncated staphylococcal nuclease is compact but disordered. *Proc Natl Acad Sci U S A* **1992**, *89*, 748-752.
(19) Metrick, C. M.; Koenigsberg, A. L.; Heldwein, E. E. Conserved Outer Tegument Component UL11 from Herpes Simplex Virus 1 Is an Intrinsically Disordered, RNA-Binding Protein. *mBio* **2020**, *11*, e00810-00820.
(20) Johnson, C. L.; Solovyova, A. S.; Hecht, O.; Macdonald, C.; Waller, H.; Grossmann, J. G.; Moore, G. R.; Lakey, J. H. The Two-State Prehensile Tail of the Antibacterial Toxin Colicin N. *Biophys J* **2017**, *113*, 1673-1684.
(21) Fagerberg, E.; Mansson, L. K.; Lenton, S.; Skepo, M. The Effects of Chain Length on the Structural Properties of Intrinsically Disordered Proteins in Concentrated Solutions. *J Phys Chem B* **2020**, *124*, 11843-11853.

(22) Paz, A.; Zeev-Ben-Mordehai, T.; Lundqvist, M.; Sherman, E.; Mylonas, E.; Weiner, L.; Haran, G.; Svergun, D. I.; Mulder, F. A.; Sussman, J. L.; et al. Biophysical characterization of the unstructured cytoplasmic domain of the human neuronal adhesion protein neuroligin 3. *Biophys J* **2008**, *95*, 1928-1944.
(23) De Biasio, A.; Ibanez de Opakua, A.; Cordeiro, T. N.; Villate, M.; Merino, N.; Sibille, N.; Lelli, M.; Diercks, T.; Bernado, P.; Blanco, F. J. p15PAF is an intrinsically disordered protein with nonrandom structural preferences at sites of interaction with other proteins. *Biophys J* **2014**, *106*, 865-874.
(24) Bremer, A.; Farag, M.; Borcherds, W. M.; Peran, I.; Martin, E. W.; Pappu, R. V.; Mittag, T. Deciphering how naturally occurring sequence features impact the phase behaviours of disordered prion-like domains. *Nat Chem* **2022**, *14*, 196-207.
(25) Plumridge, A.; Andresen, K.; Pollack, L. Visualizing Disordered Single-Stranded RNA: Connecting Sequence, Structure, and Electrostatics. *J. Am. Chem. Soc.* **2020**, *142*, 109-119.
(26) Chen, H.; Meisburger, S. P.; Pabit, S. A.; Sutton, J. L.; Webb, W. W.; Pollack, L. Ionic strength-dependent persistence lengths of single-stranded RNA and DNA. *Proc Natl Acad Sci U S A* **2012**, *109*, 799-804.
(27) Kaur, T.; Raju, M.; Alshareedah, I.; Davis, R. B.; Potoyan, D. A.; Banerjee, P. R. Sequence-encoded and composition-dependent protein-RNA interactions control multiphasic condensate morphologies. *Nat. Commun.* **2021**, *12*, 872.
(28) Elbaum-Garfinkle, S.; Kim, Y.; Szczepaniak, K.; Chen, C. C. H.; Eckmann, C. R.; Myong, S.; Brangwynne, C. P. The disordered P granule protein LAF-1 drives phase separation into droplets with tunable viscosity and dynamics. *Proc. Natl. Acad. Sci. U.S.A.* **2015**, *112*, 7189-7194.
(29) Ambadipudi, S.; Biernat, J.; Riedel, D.; Mandelkow, E.; Zweckstetter, M. Liquid-liquid phase separation of the microtubule-binding repeats of the Alzheimer-related protein Tau. *Nat. Commun.* **2017**, *8*, 275.
(30) Brady, J. P.; Farber, P. J.; Sekhar, A.; Lin, Y. H.; Huang, R.; Bah, A.; Nott, T. J.; Chan, H. S.; Baldwin, A. J.; Forman-Kay, J. D.; et al. Structural and hydrodynamic properties of an intrinsically disordered region of a germ cell-specific protein on phase separation. *Proc Natl Acad Sci U S A* **2017**, *114*, E8194-E8203.
(31) Regy, R. M.; Thompson, J.; Kim, Y. C.; Mittal, J. Improved coarse-grained model for studying sequence dependent phase separation of disordered proteins. *Protein Sci* **2021**, *30*, 1371-1379.
(32) Dannenhoffer-Lafage, T.; Best, R. B. A Data-Driven Hydrophobicity Scale for Predicting Liquid-Liquid Phase Separation of Proteins. *J. Phys. Chem. B* **2021**, *125*, 4046-4056.
(33) Bai, Q.; Zhang, Q.; Jing, H.; Chen, J.; Liang, D. Liquid-Liquid Phase Separation of Peptide/Oligonucleotide Complexes in Crowded Macromolecular Media. *J. Phys. Chem. B* **2021**, *125*, 49-57.
(34) Fisher, R. S.; Elbaum-Garfinkle, S. Tunable multiphase dynamics of arginine and lysine liquid condensates. *Nat. Commun.* **2020**, *11*, 4628.
(35) Alshareedah, I.; Kaur, T.; Ngo, J.; Seppala, H.; Kounatse, L. D.; Wang, W.; Moosa, M. M.; Banerjee, P. R. Interplay between Short-Range Attraction and Long-Range Repulsion Controls Reentrant Liquid Condensation of Ribonucleoprotein-RNA Complexes. *J Am Chem Soc* **2019**, *141*, 14593-14602.
(36) Tesei, G.; Schulze, T. K.; Crehuet, R.; Lindorff-Larsen, K. Accurate model of liquid-liquid phase behavior of intrinsically disordered proteins from optimization of single-chain properties. *Proc Natl Acad Sci U S A* **2021**, *118*, e2111696118.
(37) Dignon, G. L.; Zheng, W.; Kim, Y. C.; Best, R. B.; Mittal, J. Sequence determinants of protein phase behavior from a coarse-grained model. *PLoS Comput Biol* **2018**, *14*, e1005941.
(38) Dannenhoffer-Lafage, T.; Best, R. B. A Data-Driven Hydrophobicity Scale for Predicting Liquid-Liquid Phase Separation of Proteins. *J Phys Chem B* **2021**, *125*, 4046-4056.
(39) Regy, R. M.; Dignon, G. L.; Zheng, W.; Kim, Y. C.; Mittal, J. Sequence dependent phase separation of protein-polynucleotide mixtures elucidated using molecular simulations. *Nucleic Acids Res* **2020**, *48*, 12593-12603.
